# In a pathophysiologic state due to SARS-CoV-2, viscosity and cholesterol are two-edged swords

**DOI:** 10.1101/2025.02.17.638432

**Authors:** Ikechukwu Iloh Udema

## Abstract

Much attention has been paid to the genetic composition and molecular biology of viral particles, infection, and micro-anatomical impacts that culminate in fatalities; vaccines have been in continuous development and production; less attention is paid to the fundamental issues of thermodynamics and activation energy characterization of viral RNA replication and cell death. The study aimed at deriving equations that can be fitted to both theoretically and empirically derived data for the quantitation of other thermodynamic parameters and its cognate dimensionless equilibrium constant; some of the derived equations addressed the issue of viscosity and the concentration of cholesterol in particular as they affect translational velocity needed for the delivery of biomolecules to the site of need. The instantaneous velocities before terminal velocity are: ∼ 0.046674 m/s (cytosol); 0.141837 m/s (water). The terminal velocities were approximately equal to 3.548614 nm/s for the cytosol and 99.590626 nm/s for the water; these values were computed using literature values of translational diffusion coefficients (Di) of glucose in cytoplasm and in water. The value in water is higher than in the cytosol because of higher cytosolic viscosity than aqueous viscosity. These support the view that cholesterol and viscosity have a dual-edged effect on the pathophysiologic state orchestrated by SARS-CoV-2; higher viscosities in the membrane and in the cytoplasm enhance binding and infection and can diminish the progress of infection, respectively. Higher feasibility and rates were observed at lower thermodynamic temperatures than at higher ones, according to the outcome of the analysis of the viral binding free energy and activation energy, respectively. The dimensionless constant values were higher at the earlier time of the infection and decreased with time, exhibiting a power law relationship. It is advised, among others, that pharmaceuticals (including airborne surfactants) and drugs in solution be given at temperatures above body temperature. Swab testing should be performed on a regular basis to detect significant infections early. Future in vitro and in vivo studies on viral infection might focus on various time periods at different temperatures, above and below body temperature.

**GRAPHICAL ABSTRACT:** 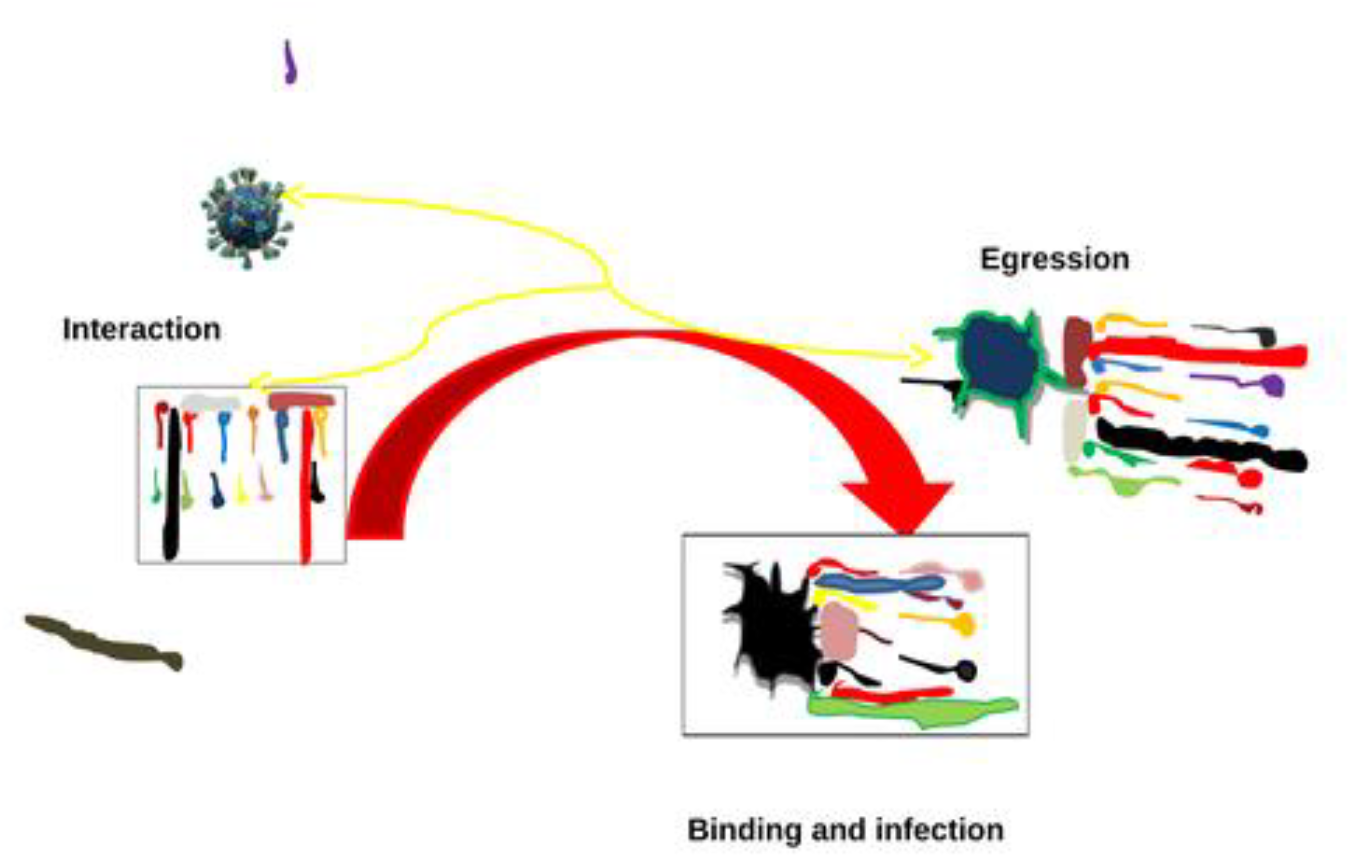

With vaccine and/or drug the viral infection can regress; without treatment there could be infection and progression into disease state; discontinuation of treatment can cause a reinfection.

## 1.0 INTRODUCTION

> *“Intellectual superiority complex and its opposite, which has yet to prevent death due to COVID-19, oppose the knowledge that no one is a tabula rasa”*.

Humanity has not yet fully overcome the death caused by COVID-19, despite the belief in an intellectual superiority complex and it’s opposite. This kind of situation might have given rise to the remark that “all theories are useful, but not all theories are valid” (Srinivasan, 2022), which was based on a theory’s apparent fault, but the author did not go so far as to criticize it due to the personality who advanced the idea. This is an example of the effect of the inferiority complex. This study has no room for such a scenario. Now, one dares not put to question the following: 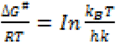, and 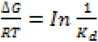 ; the former refers to the ratio of activation free energy (Δ*G*^#^) to the thermal energy (*RT*) and the latter is the case for substrate binding free energy (Δ*G*) (Sousa, *et al*., 2020); *R, T, k*_*B*_, *h, k, and K*_*d*_ are the universal gas constant, thermodynamic temperature, Boltzmann constant, Planck constant, first-order rate constant, and ligand-receptor complex dissociation constant (in unit of either mass or molar concentration), respectively. The ratio of free energy change of any kind to thermal energy is clearly dimensionless. The natural logarithm of another ratio of energies *k*_*B*_*T/hk* is also dimensionless. Therefore, no one should be blind-folded to the reality that *K*_*d*_ is either in the unit of g/L or mol. /L. In order not to confuse I with l (the lower case of L), L is used as the symbol of a liter. There is no question of either standard unit mass concentration or unit molar concentration. The standard unit concentration does not originate from experimental data. To accurately characterize the thermodynamics of virus-receptor contact and the activation-free energy associated with it, this work takes these and other factors into account. This also covers all metabolic processes that are both directly and indirectly related to viral infection, its RNA replication, and cell death.

If there isn’t any motion at the molecular, cellular, and organismal levels, life might not exist forever. However, the “creation of life” requires motion to stop. In accordance with Schrödinger’s theory, Pierce thus asserts that life is essentially a process in which matter is gathered, rearranged, and organized using energy in response to information encoded in aperiodic molecules. What appears to be a definition of life is criticized for lacking a clear mechanism for the management and reorganization of matter (Pierce, 2023). Genetics, molecular biology, developmental biology, biochemistry, and other fields might contain hints. The fundamental issue is a downward trend in entropy, or the entropy of biological systems criticized for violating the second law of thermodynamics (*i*.*e*., that entropy in a system always increases) and purported to be devoid of any explanation. “A layman’s view in this regard is that as long as there is one universe in which there are different systems, any change in characteristics in any of the systems should create opposing change either in the same system composed of subsystems or in another system” (*in the book manuscript submitted for evaluation*).

The bone of contention remains diffusional motion which may be in lateral and transverse direction both of which can affect the integrity of the membrane. Diffusion is the driving force behind passive transport (Sugano, *et al*., 2010). As such physicochemical information on the diffusion of any solute is useful for analyzing food, metabolite, hormone, enzyme, and drug delivery systems and pharmacokinetics. As such, the distributions or diffusion velocities of central nervous system drugs in the brain after they penetrate the blood-brain barrier can, also, be investigated in a manner that can be enabled by such information (Nicholson, 2015). The site of biosynthesis needed for viral replication processes can be starved of metabolites if the velocity of delivery is retarded.

In normal health, most molecules must be transported passively by diffusion: diffusion of oxygen into the blood and carbon dioxide out of the blood across the alveolar-capillary barrier; diffusion of digested food and other biomolecules; and xenobiotics, some of which may be prescription drugs and alcohol. Because of the importance of diffusion, literature is replete with pieces of information about both theoretical and experimental results and conclusions. Some theoretical investigations are intended to generate models or rather procedures that do not require special apparatus that can speed up the determination of diffusion rate and diffusivity or diffusion coefficient; in other words, computational approaches such as molecular modeling are becoming more and more desirable (Miyamoto and Shimono, 2020). This is not just about diffusion- and non-diffusion-controlled enzyme-catalyzed reactions but that the metabolites have to be transported either passively by diffusion or by active transport, neither of which can evade the nature of the medium.

In most, if not all, there are emphases on the effect of the solvent medium as per the viscosity rather than the viscosity of either the homogeneous or heterogeneous solution. The biological milieu, be it the cytoplasm, blood, *etc*., is nevertheless a heterogeneous mixture. Indeed, the plasma membrane (PM), nuclear membrane, and membranes of organelles are also heterogeneous in composition. One of the functions of PM and other membranes is selective control of molecules, supramolecules, *etc*. However, xenobiotics such as prescription drugs and alcohol find their way into the cytosol, thereby giving the impression that the selective barrier function of PM does not possess any mechanism of detecting what can be considered as original substances bearing biological relevance to the cell. Viral particles and poison can find their way into the cell. Therefore, the main secondary defense available to the living things, the higher organism in particular, is the immune system. It has been established, however, that the viral agents overthrow the host immune system in order to survive. But if these pathogens meet resistance within the cytoplasm, the potential of the viral agent to torpedo the defense mechanism can be attenuated to zero.

Be it as it may, the selective barrier function is indispensible even if it is known to provide anchorage, a function of a supporting platform or scaffolding, to pathogenic viral agents as a first step in the pathway of pathogenesis and disease outcome. Here, the coronavirus is explored as an example for the demonstration of the role of the micro-biotic environment of the cellular body that has the potential to enhance its own evasion and also therapeutic-dependent protection. In this regard, the coronaviruses are described as enveloped viruses with non-segmented, single-stranded positive sense RNA genomes (the kind that acts as messenger RNA) (Drosten, *et al*., 2003). Rahimi, *et al*. (2022) identified human coronavirus OC43, human coronavirus NL63, human coronavirus 229E, SARS coronavirus, human coronavirus HKU 1, and Middle East respiratory syndrome coronavirus as those that infest man. Among these, SARS coronavirus 1 and 2, and Middle East respiratory syndrome coronavirus are the most pathogenic (Rahimi, *et al*., 2022). The SARS-CoV-2, the choice in this investigation belongs, according to Rahimi, *et al* (2022). and the references therein, to the genus, *Betacoronavirus*.

A number of mathematical models have been created to provide guidance to health care providers and health policy makers in the development of methods for reducing the spread and virulence of SARS-CoV-2 as well as the determination of the best practices for treatment and prevention (de Pillis, *et al*., 2023). However, the models focus on modeling epidemiological dynamics and answering questions about population-level effects of interventions such as vaccination and treatment (Diagne, *et al*., 2021, Swan *et al*., 2021), while other investigators (de Pillis, *et al*., 2023) are addressing questions about protective immunity within a vaccinated individual. The models for the binding of viral particles on the membranes cannot be precluded, but the factors that enable the fixation of the viral particle are of great concern. One of such factors includes the composition of the membrane. Lipids such as phosphatidylcholine and sphingomyelin (in the outer membrane); phosphatidylethanolamine, phosphatidylserine, and phosphatidylinositol; variable amounts of cholesterol; lipid-anchored proteins confined to only one side of the membrane; integral proteins; and peripheral proteins are such examples. In fact, **t**he lipid composition of the cell plasma membrane contains 26% phosphatidylcholine, 24% sphingomyelin, and 12% glycosphingolipids. There is also the presence of lipid rafts, specialized microdomains within the cell membrane, which are composed of saturated phospholipids, sphingolipids, glycolipids, cholesterol, lipidated proteins, and glycosyl phosphatidyl inositol-anchored proteins (Simons and Ehehalt, 2002).

It is obvious that the first point of contact remains the membrane, which is the platform that can stabilize the virus preparatory to entry into the cytoplasm. The platforms in microdomains referred to as lipid rafts and associated proteins and receptors contain cholesterol apart from other lipids, which renders such domains more rigid or viscous. The cytoplasm also contains proteins, lipids, and fats, including in particular cholesterol and its derivatives, all of which contribute to the viscosity of the medium. While the level of viscosity at the membrane level enhances viral infection, high viscosity due to cholesterol, proteins, *etc*. in the cytoplasm inhibits the replication of viral materials. High viscosity can retard the diffusion of metabolites needed for the synthesis of viral RNA and scaffolding materials such as cholesterol, which are transported to membranes by HDL. Therefore, the proposition that in a pathophysiologic state due to SARS-CoV-2, viscosity and cholesterol are two-edged swords cannot be out of the question. This attribute can be explored in efforts aimed at halting viral infection. Meanwhile, much attention has been paid on genetic composition and molecular biology of viral particles, infection, micro-anatomical impact that culminate into fatalities; vaccines has been in continuous development and production; less attention is paid to the fundamental issues of thermodynamics and activation energy characterization of viral RNA replication and cell death. This study is therefore aimed at defining apparent thermodynamic and activation energy model equations that can enable the quantification of opposing energies that either drive or oppose the biosynthetic pathway that supports virulence and pathogenesis itself. Secondly, the study aims to explore the models that describe the effect of viscosity in different compartments. Other models which can be fitted to experimental or calculated data are explored for the determination of time before every events culminating to cell death. Thirdly, the impact of temperature on the progress of infection is part of the goals of the study. Fourthly, the means of achieving the goal of halting the infection and replication of viral materials are proposed.

The impact of Severe Acute Respiratory Syndrome-CoV-2 (SARS-CoV-2) assumed a catastrophic outcome of biological warfare second to nuclear war in its “bad and ugly side”. COVID-19, a result of the SARS-CoV-2 infection, is well reported as an extremely severe respiratory illness (Yu *et al*., 2023) that has claimed millions of lives worldwide by causing a high degree of hypoxia (Makanya, *et al*., 2013). Highly advanced, leading scientists should indeed take the lead in finding faults and gaps in existing models where applicable, even as it appears there are emerging new models, to enhance the realizable hope of defeating the common enemy.

## 2. BRIEF THEORETICAL PRESENTATION

### 2.1. Effective Kinetic energy

There is an understanding that viscosity and any metabolite that can either increase or decrease it, could serve to regulate influx and efflux of biomolecules both micro-and macro-compounds in out the cell membrane or the entire cytoplasm but not limited to them. The issues remain either the downward or upward regulation of diffusional processes. To this end the equations (a book under preparation) that address the effective kinetic energies in solution is given as Eq. (1) below. Equation (1) implies that 3 *k*_*B*_*T*/2 is not the effective average kinetic energy. If not, it is equivalent to stating that a fast-moving sports car can maintain its starting velocity without experiencing a reduction when it unexpectedly confronts a powerful tornado in the opposite direction. This is as absurd as the idea that the effective root mean squared velocity is equal to (3*k*_*B*_*T*/*m*_*i*_)^**½**^. Diffusing solutes encounter resistance against their motion, which ends in a terminal velocity, not only in solutions with electric field gradients. This also holds true for any type of solute that diffuses along a concentration gradient until it reaches uniform concentration or its specific destination. The viscosity of the solution (or solvent in the case of much diluted solutions) is the cause of this resistance. Higher viscosity is caused by a variety of viscogenes, including large molecular mass proteins and cholesterol (and other saturated fats). Although high viscosity can obstruct the metabolic pathway necessary for the synthesis of cholesterol and other materials needed for the virus’s replication, it can also increase the infectivity of pathogens like the coronavirus by giving them a platform to contact the cell membrane. A book is being prepared that addresses other problems related to Equation (1) (Udema, 2016).

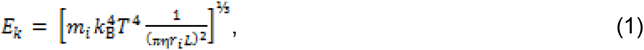

where *m*_i_, *k*_B_, *T*, η, *L*, and *r*_i_ are the mass of any solute, Boltzmann constant, thermodynamic temperature, viscosity of the medium rather than just the solvent alone, the cube root of the molar volume (*m*_1_/ρ_1_)^⅓^, where *m*_1_ and ρ_1_ are the mass of a molecule of water and density of pure water at the prevailing temperature, and the hydrodynamic radius of the solute, respectively; *E*_k_ is the thermally influenced translational energy. Note that the medium stated earlier is an inhomogeneous body composed of fluid, solutes of different sizes, intracellular bodies such as organelles, *etc*. Given that the translational diffusion coefficient (*D*_i_), which should be determined under the influence of the medium (this also includes the biological membranes), is according to the Einstein and Stoke equation, given as *k*_*B*_*T*/6πη*r*_i_. So πη*r* can be expressed as *k*_*B*_*T*/6*D*_*i*_. Substitution into Eq. (1) gives alternative to Eq. (1) represented by Eq. (2) below:

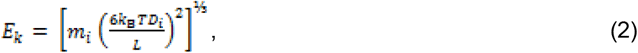

### 2.2. Misuse and misrepresentation of Einstein’s diffusion equation

While Einstein–Smoluchowski equation of diffusion has assumed different forms, the most notable of all is the following equation:

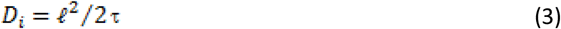

where ℓ and τ are the root mean square displacement of the solute and its corresponding duration in seconds. While τ can be chosen arbitrarily for whatever reason, and if *D*_*i*_ is known *ab initio*, the expected value of ℓ can be calculated given well-defined conditions such as *T* and η_i_ of the medium. It amounts to misapplication and misrepresentation of Eq. (5) if, given *D*_*i*_, one chooses to quantify τ given an arbitrarily chosen value of ℓ, such as the long dimension length of the intracellular medium. The ℓ is strictly a root mean square displacement in line with the original Einstein’s formalism. In other words, only *D*_*i*_ and length can be determined experimentally. The parameter, ℓ, is an outcome of statistical treatment, though no one has shown how one can sum up the square of each displacement of every molecule in trillions of them and above in a way that a layman can understand. The mean square displacement (MSD) and root MSD have no alternative interpretations that are different from the view of a well-respected Jewish American scientist, Albert Einstein. “While “anti-author” may often lead to the rejection of novel ideas by the author, anti-idea does not always translate to anti-author”. Thus, there could be other transformations that can culminate in Eq. (3) or a direct transformation of the latter into other forms for possible application in a different setting.

However, as was previously established, there are always values of ℓ that correspond to values of τ if Di is known for a particular medium whose physiochemical parameters are known. To specify the values of τ and ℓ for any solute or particle in general, information about *D*_*i*_ alone is insufficient.

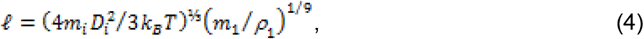

Equation (14b) proposes that the magnitude of ℓ is inversely proportional to the cube root of the temperature against the known fact that a longer displacement may show a linear relationship with the temperature, although not the cube root of the temperature, the presentation in Eq. (14b) notwithstanding.

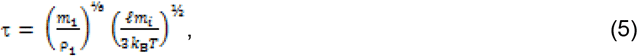

Earlier in the text, the characteristic of the medium is taken into account in the consideration for the values of η and *D*_i_. Hence, the issue of anomalous diffusion becomes penitent because such is due to the effect of heterogeneous and asymmetric medium offered by biological membranes and all media enveloped by membranes. Anomalous diffusion, the departure of the spreading dynamics of diffusing particles from the traditional law of Brownian motion, is regarded as a signature feature of a large number of complex soft matter and biological systems (Sposini, *et al*., 2022). In general, anomalous diffusion processes are those where the mean square displacement does not conform to the linearity expected in Eq. (5), and as may be applicable to another version of the latter given as (Oliveira *et al*., 2019):

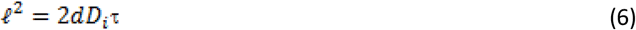

where *d* is the spatial dimension. In other words, there is a non-linear relationship between squared displacement and time. Anomalous diffusion includes sub- and super-diffusion; the subdiffusion aspect is of interest in this investigation. The fractional Brownian motion (fBm) model faithfully describes the diffusion of particles in a viscoelastic fluid (Ernst, *et al*., 2012), and it has often been argued that molecular crowding in the cell gives rise to microviscosity and therefore to anomalous diffusion (Woringer, *et al*., 2020). This presentation implies that there could be the occurrence of normal diffusion and at least subdiffusion in the same medium. This is confirmed by the report that normal and abnormal regimes can coexist in proportions that depend upon the cellular organization (Kalwarczyk, *et al*., 2012, Dix and Verkman, 2008). In addition, the measured diffusion coefficients are observed to decrease rapidly as the crowder size increases, not only as a result of the space occlusion but also because of biopolymer distribution and interactions (the latter including solvent-mediated correlations or hydrodynamic interactions) (Elowitz, *et al*., 1999, Wang, *et al*., 2011, Ando and Skolnick, 2010).

Anomalous diffusion processes that do not agree with the linearity expected by Eq. (5) remain relevant to this study, while Brownian diffusion cannot explain the physics of disordered systems, as exemplified by the composition of the cytoplasm, for instance. Interestingly, a ubiquitous observation in cell biology is that the diffusive motion of macromolecules and organelles is anomalous, *i*.*e*., the MSD change with time is typically characterized by a sublinear increase. In most instances, this sublinear increase of the MSD with time can be fitted to a power-law relation with an exponent α < 1, which justifies the term “subdiffusion.” Subdiffusion is usually attributed to cellular crowding, spatial heterogeneity (or inhomogeneity), or molecular interactions (Woringer, *et al*., 2020). The fractional Brownian motion (fBm) is a Gaussian stochastic process characterized by a zero mean value and the covariance function given as:

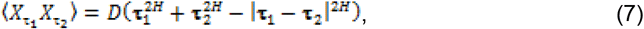

where *D* is a proportionality factor with physical units of ℓ^2^/**τ**^2*H*^ commonly referred to as generalized diffusion coefficient and *H* ∈ (0, 1) is the traditionally used Hurst index, such that the anomalous diffusion exponent is α = 2*H* (Woringer, *et al*., 2020). Elsewhere, however, 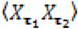 is regarded as a correlation function for **τ**_1_ > **τ**_2_ > 0 (Trovato and Tozzini, 2014). Nonetheless, both positions border on higher-level statistical issues that may not bear direct relevance to the current study. Thus, fBm describes a process that can be subdiffusive (*H* < ½), diffusive (*H* = ½), or superdiffusive (*H* > ½) (Sposini, *et al*., 2023).

Two quantities that can be extracted from the MSD are the apparent diffusion coefficient *via* the relationship *D*_i_ = MSD/6**τ** and the anomalous exponent α from the slope of the MSD in the log-log plot. The latter equation is different from Eq. (3), but it is not very apparent why there is a preference for its use in the literature. Be it as it may, either equation being used suggests that nonlinearity takes the backseat since any determination of the viscosity coefficient cannot be done without the equations. This is especially true if a state function is used in a situation that should be handled by a path function since the main focus is on the start of diffusional motion, regardless of any resistance, cul-de-sacs, transient interactions, or other impediments that lengthen the time it takes to reach the intended destination.

In packed media, such as the cytoplasm and any membrane, randomness is less likely to occur. The diffusivity can be computed given the transit time and a specific thickness and distance between two cytosolic locations. The value *D*_i_ is then combined with the corresponding diffusion coefficient in dilute solution, *D*_0_, to obtain the quantity, ln(*D*_0_/*D*_i_) = ln(η_i_/η_0_); the latter measures the increase of apparent solution viscosity η_i_ relative to the one in dilute conditions, η_0_ (Trovato and Tozzini, 2014). The scenario presented herein is that given chosen time, the MSD must be experimentally determined. In conducting this study, there is awareness that biological media are entirely different from ordinary dilute and even concentrated aqueous solutions. Though the unit of life is the cell, it is still under the influence of systems such as the nervous and hormonal systems in particular. The cell, in turn, is a microsystem represented by organelles within the complex and viscous fluid medium. Thus, against this backdrop, there is the phenomenon of anomalous diffusion characteristic of biological media due mainly to very high viscosity and highly crowded medium.

### 2.3. Viscosity

One parameter that describes cytoplasmic rheology is “fluid-phase viscosity,” defined as the microviscosity sensed by a small solute in the absence of interactions with macromolecules and organelles (Fushimi, 1991). Popular pictorial representations of the aqueous environment within cells suggest that the crowding might seriously hinder solute diffusion (Fulton, 1982)—a major determinant of metabolism (Welch and Easterby, 1994), transport phenomena, signaling, and cell motility. Besides, there is support for the view that the cytoplasm is more like a bag of slightly viscous water than a complex gelatinous mass, at least with respect to the effect on solute diffusion (Seksek, *et al*., 1997). This seems to go against the finding that the solvent viscosity of cytoplasm was not significantly different from that of water and showed no spatial variation (Luby-Phelps, e*t al*., 1993).

The microviscosity of a solution is the resistance to motion experienced by a molecule in the solution. This contrasts with the macroviscosity measured by conventional viscometers, which is a bulk property of the solution (Copeland, 2002). This is a point of contention in this study, notwithstanding the fact that any solute subject to microviscosity cannot be in isolation from the prevailing macroviscosity of the solution. Because increases in microviscosity increase the resistance to molecular motions in solution, the rate of diffusion is slowed down (Copeland, 2002). Polyethylene glycol, like any other polymeric viscogenes, influences the macroviscosity only, while monomeric viscogens, such as sucrose and glycerol, affect both the macro- and micro-viscosities. Hence, adding monomeric viscogenes into a solution or fluid medium in general could be the best and simplest way of increasing the microviscosity (Copeland, 2002). A takeaway hypothesis in this case is that both micro-and macro-viscosities have the potential to slow down intracellular interaction of species, considering the impression given by Eq. (26b). Implied is also a decrease in the stability of the viral particles on the membrane, whose lipid raft may have been made more fluid due to the exit of cholesterol in particular and some receptor proteins. These scenarios can be modeled first as applicable to the cytoplasm, where metabolism essential for the replication of viral particles occurs. For any of the reversible biochemical reactions in the cytoplasm, an equation similar to that observed in the literature (Copeland, 2002) is explored. This is premised on the view that if virulence is reduced due to heightened viscosity due to the increasing presence of viscogens, cholesterol, monomers, such as nucleic acids, glycerol, fatty acids, simple sugars, *etc*., and polymers of whatever kind, then viscosity gives the impression that the rate of infection may be diffusion dependent considering the metabolome that supports the proliferation of the virus.

Thus,

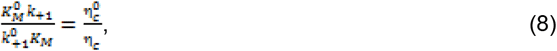

where *k*_+1_ and 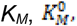 and 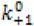are the first-order catalytic rate (often found as *k*_*cat*_ in the literature) and Michaelis-Menten constant with viscogenes, Michaelis-Menten constant and first-order catalytic rate without viscogenes; 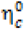 and η_c_ are viscosities without and with viscogenes respectively. Equation (30) is similar to the equation [In *D*_0_/*D*_i_ = Inη/η_0_ (Trovato and Tozzini, 2014). It can be seen straight away that in Eq. (30), the specificity constant is reasonably inversely proportional to the viscosity coefficient. A straightforward inference is that the viscosity coefficient is directly proportional to the *K*_*M*_ and, consequently, to the *K*_*d*_ (equilibrium dissociation constant), which should impact the rate at which any type of *LR* (ligand receptor complex) or *ES* (enzyme-substrate complex) forms.

### 2.4. Preemptive thermodynamics and activation energy consideration pertinent to the binding and replication of viral RNA and cell death at the linear phase

The fundamental idea behind the proactive study of thermodynamics and activation energy (Arrhenius and Gibbs activation energies) is that the coronavirus’s binding spike protein’s Gibbs free energy equation no longer appropriately uses *K*_*M*_ or *K*_*d*_. It is only because of its historical background. To put it another way, it is consistent with the educational principle of starting with the known—whether true or false—and working your way up to what appears to be previously unknown. Be it as it may, the notion that *K*_*M*_ may stand for equilibrium constant (“this may remain acceptable in advancing and highly advanced scientific communities and perhaps some undeveloped countries where I belong”) is repudiated given the following: Under specified conditions, it is considered that it is the ratio, *K*_*M*_ = (*k*_−1_+*k*_*+*1_)/*k*_*f*_ ≈ *k*_−1_/*k*_f_ (if *k*_*+*1_≪ *k*_*+*1_), that matters, and *K*_*M*_ can be estimated experimentally, though, often with outliers. Simply put, *K*_*M*_ is not a thermodynamic equilibrium constant; it is rather an approach to the zero-order constant, because it specifies the substrate concentration at which half the maximum velocity of the catalytic action is done.

While an approximation such as *k*_−1_/*k*_*f*_ (subscripts −1 and *f* stand respectively for backward first-order rate constant and forward second-order rate constant) should be dissociation constant, the inverse should be an association constant or equilibrium constant in principle. The concern as expressed elsewhere (Udema and Onigbinde, 2019) is that the thermodynamic equilibrium constant (*K*_*eq*_) for the determination of apparent Gibbs free energy of any process must be dimensionless, similar to the expectation from Eyring’s equation given as (Eyring, 1935):

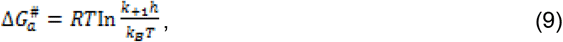

The ratio *k*_−1_*h/k*_*B*_*T* has no dimension (emphatically unitless). *K*_*M*_ or *K*_*eq*_ has the unit of mol. /L or g/L when considering unit concentration of each reactant and product; this, then calls for a unitless thermodynamic equilibrium constant.

### 2.5. Dimensionless equilibrium constant (*K*_*eq(*δ)_) based on mole fraction

It has to be made abundantly clear that it is only in an isomerization reaction that the fraction of the substrate taken up by the enzyme to form *ES* within a given duration yields the new isomer whose concentration is equal to [*ES*], which is also equal to the concentration of the fraction of substrate taken up. However, in other equilibrium reactions (reversible reactions) in which two or more products, including water, if applicable, are released, the concentrations of such products may not be equal to the concentration(s) of the substrate(s). Initial contact with a pathogen does not lead to infection of all cells of any tissue or organ, just as not all substrate molecules are converted to products in a short period, even if [*E*] is higher than [*S*]. Therefore, free substrate, free enzyme, free ligand (spike protein), and free receptor (lipid raft) should be explored in deriving mole fraction and number density fraction, as the case may be. Before proceeding, recall that the enzyme catalyzed condensation of 2 molecules of acetyl-CoA to acetoacetyl-CoA and CoA-SH. This is an example in which the number of moles of reactants and products are equal. This scenario does not require the invocation of mole fraction in the derivation of a dimensionless equilibrium constant because doing so reproduces the equation (Eq. (10)) below. In other words, *K*_*eq*_ is the same as *K*_*eq(*δ)_.

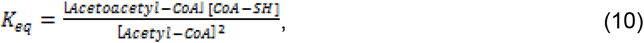

Note that *K*_*eq*_ in Eq. (10) is dimensionless. Based on Copeland (2002) formulation, the Arrhenius activation energy (*E*_a_) is related to Eyring free energy of activation as follows:

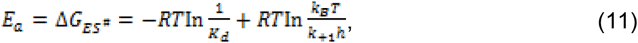

where *R* and *T* are respectively, universal gas constant and thermodynamic temperature. But *K*_*d*_ has a unit mol./l or g/l unlike *K*_*eq*_ in Eq. (10). Substitution of Eq. (10) into *RT* In*K*_*eq*_ gives:

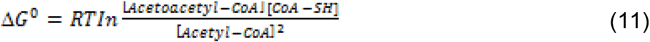

### 2.6. Mole fraction of biomolecular reactants and enzyme-substrate complex without solvent

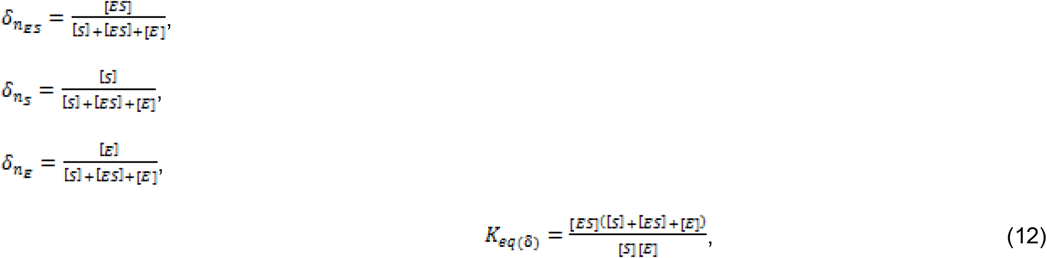

Here [*S*] and [*E*] are, respectively, the free substrate and enzyme concentrations. However, given that the *K*_*M*_ is [*E*][*S*]/[*ES*] (*i*.*e*. if free substrate concentration, [*S*] ≈ initial concentration, [*S*_0_]), than 1/*K*_*M*_ = [*ES*]/[*E*] [*S*_0_] and substituting the latter into Eq. (12) gives:

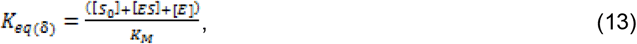

There is a need to advise that in situations where either the initial total substrate concentration ([*S*_0_]) is ≫ the total enzyme concentration [*E*_0_], a standard quasi steady-state approximation (QSSA) scenario or [*S*_0_] ≪ [*E*_0_], a reverse or total QSSA model, the following, should respectively, be taken into account:

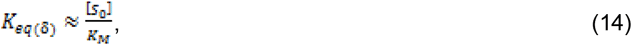

where of course, *K*_*M*_ = (*k*_-1_+*k*_+1_)/*k*_*f*_ and *k*_-1_ is first-order rate constant for the dissociation of ES.

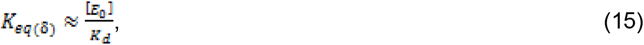

And where of course, *K*_d_ = *k*_-1_/*k*_*f*_. Nonetheless, there may be a scenario in which [*S*_0_] ≈ [*E*_0_] such that:

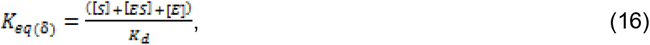

Note that in all cases, the free enzyme and free substrate are to be directly explored except on the condition that [*S*_*T*_] ≫ *K*_*M*_ and [*E*_*T*_] such that the free concentration of the substrate [*S*] ≈ [*S*_*T*_]. The same concern is applicable to ligand-receptor relationship. Recall that *K*_*d*_ = [*E*][*S*]/[*ES*], and if that is the case, the enzyme is far from being saturated. If the free substrate concentration ([*S*]) ≈ initial substrate concentration [*S*_0_], the *K*_*M*_ can be introduced in place of *K*_*d*_.

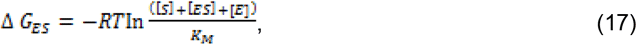

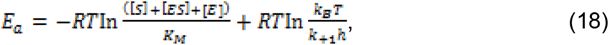

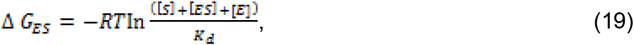

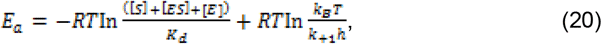

In considering the role of activation energies (Arrhenius and Gibbs free energy of activation), the Copeland (2002) model is carefully explored without giving the impression that Gibbs free energy of activation is a part of Arrhenius activation which being an empirical equation is not similar to the former which has a theoretical basis. Therefore, if the Arrhenius equation is applied to aqueous phase reactions and for the fact that 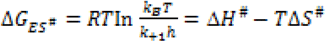 while Δ*H*^#^ = *E*_a_ - *RT* it may be reasonable to drop 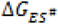 from Eqs (43), (44), and (45). Subsequent equations, where applicable, should take that into account. This position is supported by the Arrhenius equation given as: 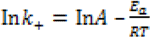 where *A* is the pre-exponential factor. Consequently, for the purpose of this investigation, such subsequent equations based on Copeland (2002) model are taken as “special cases” of activation energy equations.

### 2.7 The dimensionless equilibrium constant for infectivity, viral replication, and cell death

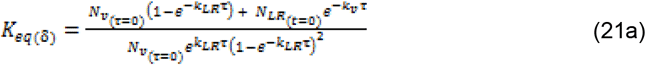

The symbols, *N*_*v*_, *N*_*LR*_, *k*_*v*_, *k*_*LR*_, and τ stand for number density of the virus, number density of lipid raft (there may be more than one lipid raft per cell membrane or cell); it is also used as the number density of the cells (Simons and Ehehalt, 2002), first-order rate constant of viral replication, first-order rate constant for cell death, and duration of events-the replication of virus and death of cells; *LR* and *v* denote lipid raft and virus respectively.

The free energy component for infectivity is:

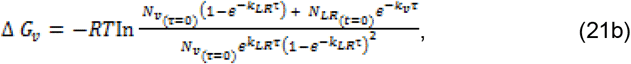

The Eyring version of activation energy that is Gibbs free energy of activation is given as:

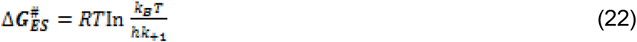

where *k*_+1_ represents either *k*_*v*_ or *k*_*LR*_.

This study focuses on SARS-CoV-2 and the molecular promoter cholesterol, whose synthesis, transport to the site of action, exit, and inhibition have either facilitated or impeded the development and progression of virulence. Again, applying Copeland’s model Copeland (2002) equation (which addresses activation energy) to the synthesis of cholesterol yields the following equation but with caveats:

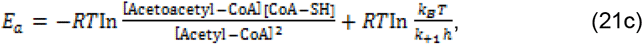

Based on Eqs (21c) and (11), one can hypothesize that the controlled inhibition of the production of AA-CoA, the controlled inhibition of the supply of A-CoA *in vivo*, and reduced consumption of cholesterol-rich food may be a means of halting the progress of the pathogenesis of SARS-CoV-2. The consumption of unsaturated fatty acids and reduced consumption of carbohydrates, which respectively take into account the needs of the cardiac muscles and the brain, may be helpful in addition to thermodynamic infeasibility and higher activation energies in halting the progress of pathogenesis. As long as a higher concentration of A-CoA is needed to synthesize AA-CoA, thermodynamic infeasibility should be the case. This is in addition to the heightened Gibbs free energy of activation made possible with a much lower *k*_+1_. Ultimately, further cell death made possible by SARS-CoV-2 infection (da Silva *et al*., 2021) can be halted as long as *k*_*LR*_ and *k*_*v*_ decrease.

There should not be doubt about the possibility of cell death orchestrated by SARS-CoV-2 in that seven TEM scans show that the density of virions within infected cells is 10^5^ per 1 pL. Such a number is hosted in the human cells, whose volume per cell is 1 pL, resulting in a cellular mass of 1 ng (Crapo *et al*., 1982, Stone, *et al*., 1992, da Silva, *et al*., 2021, Sender, *et al*., 2021) within a single infected cell at any point in time. Among any other antidotes, it may be a reasonable hypothetical proposition to deny the invading SARS-CoV-2 a stable base that enables it to inject or release viral material into the cell if and only if the cholesterol and possibly protein content of the lipid raft, estimated to be 20 protein molecules per raft (Simons and Ehehalt, 2002), is reduced. Generally speaking, the density of membrane proteins has been estimated to be around 20,000 molecules per μm^2^, giving the plasma membrane about 4 × 10^7^ protein molecules in a surface area of 2000 μm^2^. The fact that the number of 50-nm rafts is estimated to be about 10^6^ lends credence to the need to disorganize them as the sheer number promotes binding of SARS-CoV-2, whose spike protein binding sites are shielded extensively by glycosylation (Senders, *et al*., 2021).

The Δ*G*_*ES*_ and *E*_*a*_ values become more positive and higher in magnitude with higher and lower values of *K*_*M*_/*K*_*d*_ and *k*_*+*1_, respectively. *De novo* biosynthesis produces a higher concentration (≈ 700–900 mg/day) of cholesterol than exogenous sources, that is, dietary sources (300–500 mg/day). Storage, *de novo* biosynthesis, and transport to target points, *e*.*g*., the membrane, are ways of regulating cholesterol levels (Shi, *et al*., 2022). The hypothesis is that enzymes such as thiolase (for the synthesis of acetoacetyl-CoA) and HMG-CoA synthase (for the synthesis of acetoacetyl-CoA) need to be substantially inhibited. At the same time, exogenous sources should be clinically regulated so that the role of cholesterol in the infectivity of SARS-CoV-2 can be attenuated while sustaining the synthesis of steroid hormones and bile acids. Inhibition usually increases the value of the *K*_*M*_ and *K*_*d*_ such that the synthetic pathway becomes thermodynamically not feasible as the activation energies in the Arrhenius and Eryring versions become much higher than normal (Eqs (57), (58), (59) and (60)).

The processes involved in viral-host cell interaction, biomolecule trafficking in and out of the liver, uptake by the lipid raft, targeted effects on the nucleus, and vital organelles, including the mitochondria in particular, are depicted artistically in Figure 1. Tissue respiration is compromised when the integrity of the mitochondrial membrane is compromised, which lowers the amount of ATP—a molecular energy— produced. This ought to support the necessity of this study.

**Figure 1:**
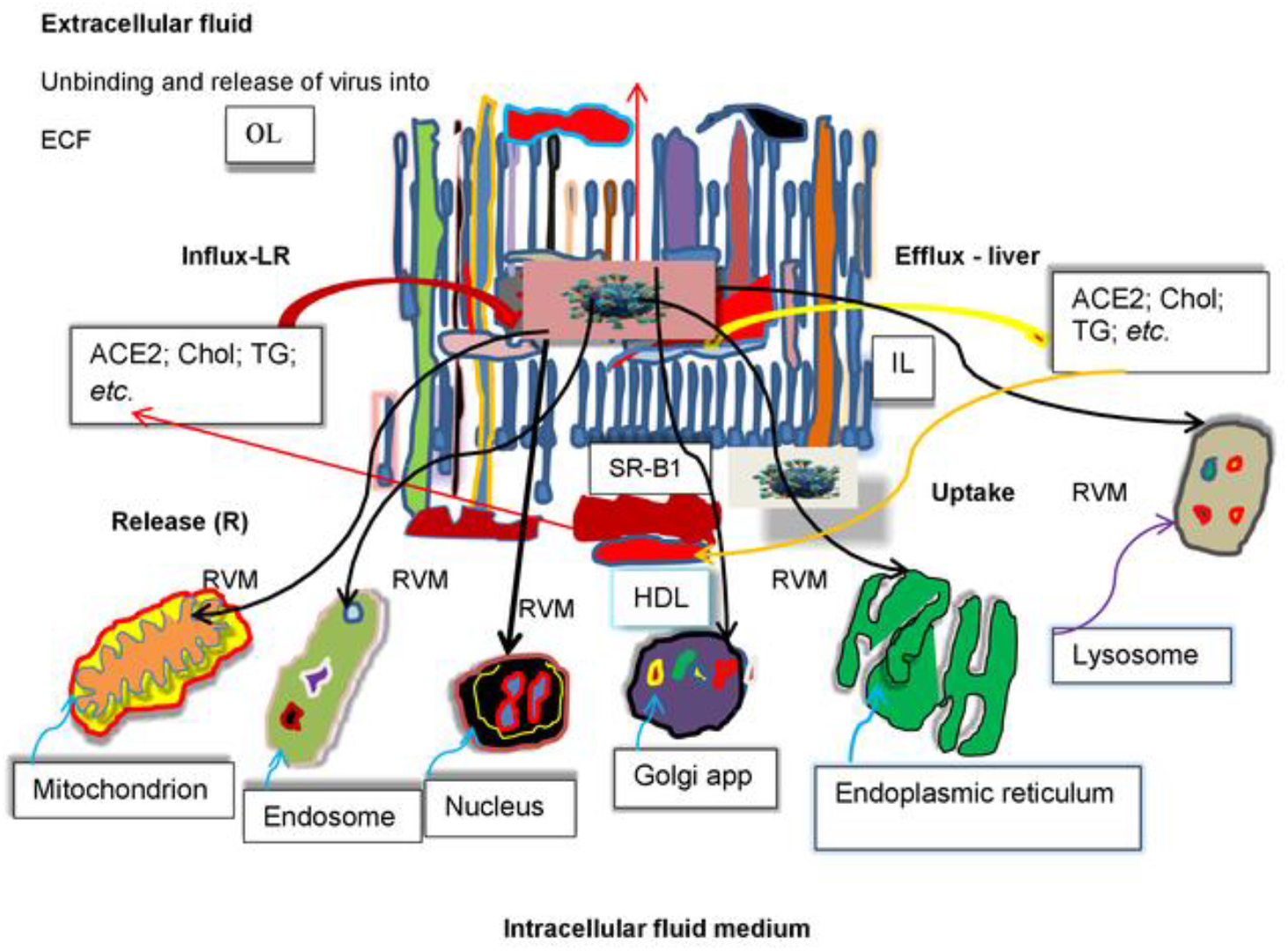
Graphics summarizing the interaction of SARS-COV-2 with the cell membrane and consequently, either infection or disinfection. RVM. LR. IL, OL. Choi, ACE2. TG. HDL. and SR-B1 denote release of viral materials, lipid raft, inner leaflet of membrane bilayer (MB), outer leaflet of MB, cholesterol, angiotensin-converting enzyme 2. tnglyceride. high-density lipoprotein, and scavenger receptor B type 1: black arrows designate targets for the impact of infection.

An experimental investigation has revealed that the presence of proteins, the spike proteins (sSpike), induces the removal of membrane lipids, both in the presence and in the absence of ACE2, suggesting that sSpike molecules strongly associate with lipids, and strip them away from the bilayer, *via* a non-specific interaction. A cooperative effect of sACE2 and sSpike on lipid extraction has also been observed (Luchini, *et al*., 2021). The presence of the protein produced a remarkable degradation of the lipid bilayer. A significant reduction in surface coverage was observed.in membranes from synthetic and natural lipids. The binding of sSpike to ACE2 triggers conformational changes that promote the fusion between the host cell membrane and the virus envelope, thereby allowing the viral RNA to be released into the host cell. Compared to other coronaviruses, the spike proteins from SARS-CoV-2 have a very low dissociation constant (14.7 nM) for the binding to ACE2, which makes SARS-CoV-2 highly infectious (Lan, *et al*., 2020). It must be stated however, that the work of Luchini, *et al*. (2021) falls short of mentioning the role of lipid rafts in which ACE2 is located, and as well as scavenger receptor B type 1 (SR-B1) which facilitates the ACE2-dependent entry of SARS-CoV-2 (Wei, *et al*., 2020)

Specifically, the equilibrium dissociation constant (*K*_*d*_) of ACE2 and SARS-CoV-2 RBD is 4.7 nM, and of ACE2 and SARS-CoV RBD is 31 nM (Lan, *et al*., 2020); other reports include ACE2 and the SARS-CoV-2 spike trimer *K*_*d*_ found to be 14.7 nM compared with that between ACE2 and SARS-CoV RBD–SD1(*K*_*d*_ of 325 nM) (Wrapp, *et al*., 2020). This large difference is speculated to be due to the different proteins used in the assay or because of other unknown reasons (Lan, *et al*., 2020). These values suggest that molar concentrations of the ligand and receptors were explored. The technology for measuring the molar concentrations of the species while with the cell membrane and virus is best known to the experimentalist. The performance of thermodynamic and activation energy analysis demands that, apart from equations derived earlier, another equation (“This could as well be described as a nascent equation” being unknown earlier) with dimensionless *K*_*eq*_ (or 1/*K*_*d*_) is derived as follows:

### 2.8. Mole fraction of viral material, the receptor-binding domain (spike protein 1), cellular biomolecule, the ACE2, and viral material-cellular biomolecule complex

This derivation is suitable if a biomolecular level involving the spike protein (designated as the ligand, *L*), the receptor (*e*.*g*., ACE2, designated as *R*, the enzyme), which is also an enzyme, and the receptor-binding domain (RBD, also taken as *R*) is the case. Thus, the dimensionless equilibrium constant (*K*_*eq*(δ)_) equation is:

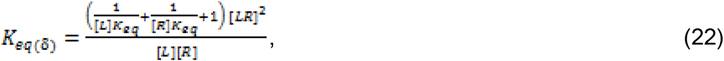

The preceding equations assume that [R] ≫ [L] as to imply absence of saturation.

The appearance of *K*_*M*_ in the equations presupposes that initial substrate concentration ([*S*_0_]) ≫ initial enzyme concentration ([*E*_0_]), and [*S*] ≈ [*S*_0_]. The concentration of all chemical species must be in mol./L. We can redirect our focus to a situation where the molar concentration of the SARS-CoV-(2) spike protein (S-C-(2) SP)-angiotensin converting enzyme (ACE) complex (S-C-(2) SP-ACE) remains uncertain or unidentified. The figure 2, in parenthesis, indicates either SARS-CoV-2 or SARS-CoV. Based on the 1:1 binding model, the relation between free [*S-C*-(2) *SP*], [*ACE*], and [*S-C*-(2) *SP-ACES*] can be stated as follows:

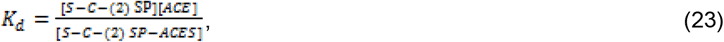

**Figure 2:**
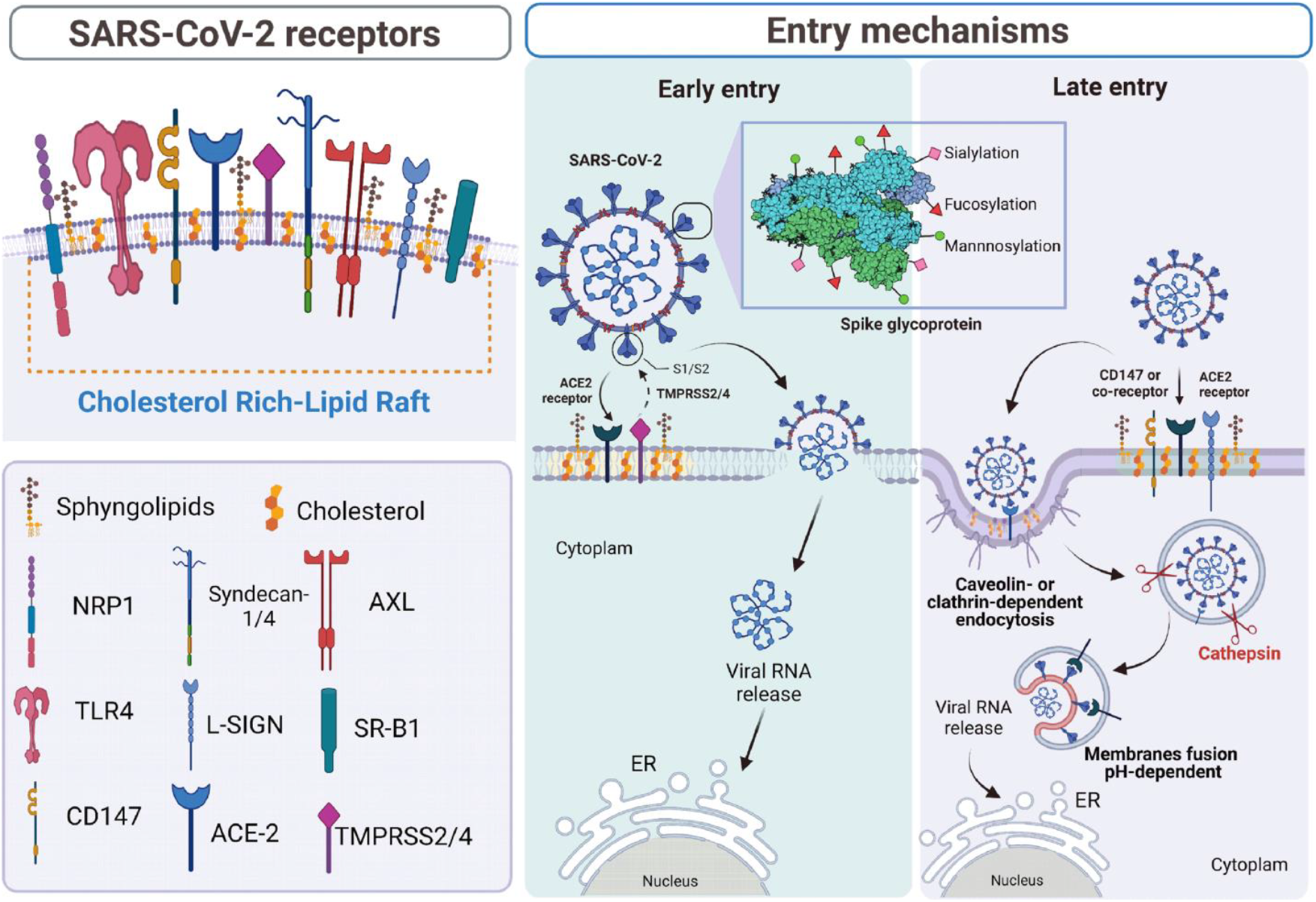
A graphic picture summarizing all processes from viral contact with cell membrane till cell death. Palacios-Rápalo, *et al*. (2021)

Moving forward, there is a need to point out that in none of the equations where *K*_*d*_ or *K*_*M*_ appears should [*ES*] or [*LR*] be > [*E*] or [*R*] and [*S*] or [*L*] for obvious reason. Based on this 1:1 model (otherwise referred to as stoichiometric ratio), [*S-C*-(2) *SP*] and [*ACE*] are equal ([*S-C*-(2) *SP*] = [*ACE*] = μ) such that,

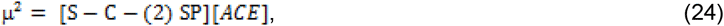

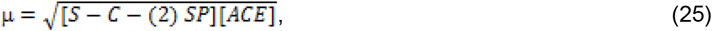

Taken into account the possibility that there are free [*ACE*] and [*S-C*-(2) *SP*] leads to the following deductions. The concentration of free ACE and S-C-(2) SP is μ − μ_C_ where μ_C_ stands for the molar concentration of the complex, [*S-C*-(2) *SP-ACE*]. This should be the case where [*ACE*] ≈ [*S-C*-(2) *SP*].

Although Eq. (24) shows the product of different biochemical species concentrations that are equal as intended, it nevertheless, gives impression that [*S-C*-(2) *SP*] and [*ACE*] may not always be equal.

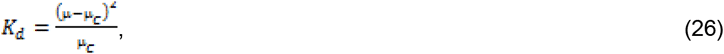

Therefore, the unknown (μ_C_), can have a quadratic solution as follows:

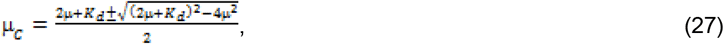

A similar equation is documented in the literature (Copeland, 2002, Schnell and Maini, 2002). It is the case in which [*L*] ≈ [*R*]. It is obvious that two positive roots, higher and lower roots, are likely outcomes of Eq. (27). Most often, *K*_*d*_, [*S-C*-(2) *SP*], and [*ACE*] are known. Be it an objective or subjective reaction for or against Eq. (27), considering more advanced equation(s) as opposed to the nascent, or preferably, *primitive* state of the equation, nevertheless, it serves as a hypothetical approach for the estimation of the SARS-CoV-ACE complex. Be it as it may, the higher root, which incidentally is higher than [*L*] and [*R*], cannot be valid considering the demands of mass conservation law. This does not foreclose its application that will shortly unfold. The mole fractions are:

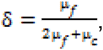

where δ is the mole fraction of free [*ACE*] and [*S-C*-(2) *SP*] which are seen to be equal; μ_*f*_ = [*ACE*] − [*S-C-* (2) *SP-ACE*] = μ−μ_*C*_.

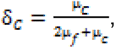

Then, *K*_*eq*(δ)_ = δ_C_/δ^2^: Substituting the equations of numerator and denominator into the latter gives the thermodynamic equilibrium constant as follows:

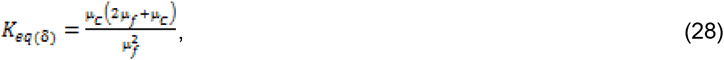

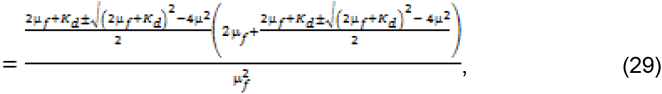

If on the other hand as stated earlier, [*S-C*-(2) *SP*] and [*ACE*] are not equal in molar terms, then, 2μ_*f*_ is replaced with μ_*SAR*_ + μ_*ACE*_, where μ_*SAR*_ and μ_*ACE*_ stand for [*S-C-*(2) *SP*] and [*ACE*] respectively; therefore, Eq. (28) is restated as:

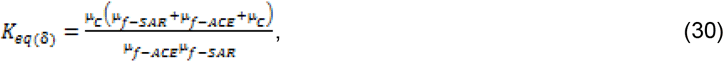

In the same vein, Eq. (79) is then transformed to:

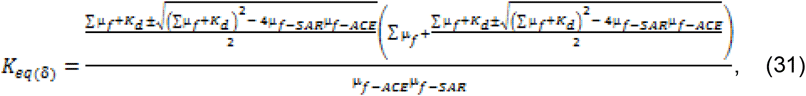

where Σμ_*f*_ = μ_*f*-*SAR*_.+ μ_*f*-*ACE*_ Note that in μ_*SAR*_ the subscript SARS spike protein (shortened to *SAR*) can be replaced with the receptor binding domain (RBD) where necessary. Besides, the free *R* and *L* (or free *E* and *S*) may be appropriate at equilibrium such that μ_c_ (or [*ES*]) should be subtracted from μ_f-ACE_ and μ_f-SAR_ (or [*ES*] should be subtracted from [*S*_0_] and [*E*_0_]). Application to the thermodynamic and activation energy questions is as illustrated earlier with 1/*K*_*d*_ (or *K*_*a*_). Equations (29) and (31) are suitable for Table 1 and Table 2 respectively.

**Table 1:** [*LR*] values at different thermodynamic temperatures plotted versus [*L*].

|  | WT | D614G |
| --- | --- | --- |
|  | **$[LR]/nM$ values** |  |
| **$T/K$** | 298.15 |  |
|  | 1.1193 | 1.1199 |
|  | 2.2285 | 2.2299 |
|  | 4.4565 | 4.4598 |
|  | 8.9209 | 8.9294 |
|  | 17.8519 | 17.8962 |
|  | 23.2530 | 23.2795 |
|  | 23.2743 | 23.2814 |
| **$T/K$** | 303.15 |  |
|  | 1.0898 | 1.1052 |
|  | 2.1670 | 2.1990 |
|  | 4.3201 | 4.3710 |
|  | 8.5722 | 8.7508 |
|  | 16.4232 | 17.1963 |
|  | 22.2219 | 22.7394 |
|  | 22.9898 | 23.1390 |
| **$T/K$** | 310.15 |  |
|  | 1.0457 | 1.0699 |
|  | 2.0754 | 2.1255 |
|  | 4.1203 | 4.2291 |
|  | 8.0888 | 8.3485 |
|  | 15.0248 | 15.7325 |
|  | 20.9567 | 21.6247 |
|  | 22.9898 | 22.7937 |

The values of $[L]/nM$ are: 1.12, 2.23, 4.46, 8.93, 17.2, 35.20, and 71.40 (Ozono, et al., 2021). The computed values of $[LR]$ are based on Eq. (27).

**Table 2:** Thermodynamic and activation parameters and cognate dimensionless equilibrium constants.

|  | WT |  |  | D614G |  |  |
| --- | --- | --- | --- | --- | --- | --- |
| **T/K** | 298.15 | 303.15 | 310.15 | 298.15 | 303.15 | 310.15 |
| **K<sub>eq</sub>($\delta$)** | 1598.93 | 41.17 | 16.84 | 23285.66 | 78.60 | 24.49 |
| **E<sub>a</sub>/e. (5)**<br>**(J/mol.)** |  | 6.65 |  |  | 7.80 |  |
| **$\Delta G$**<br>**(J/mol.)** | -18287.14 | - 9370.17 | -281.26 | -24926.90 | -11000.31 | - 8247.30 |
| **$\Delta H$/e. (5)**<br>**(J/mol.)** |  | -2.81 |  |  | -4.22 |  |
| **T$\Delta S$/e. (5)**<br>**(J/mol.)** | -2.63 | -2.72 | -2.75 | -3.97 | -4.11 | -4.15 |
| **$\Delta S$/e. (3)**<br>**(J/mol. /K)** | -0.88 | -0.90 | -0.89 | -1.33 | -1.36 | -1.34 |

The alphabet e stands for exponent. Data explored for calculations are from the literature (Ozono, *et al.*, 2021)

### 2.9. Graphical determination of *K*_*eq*(δ)_

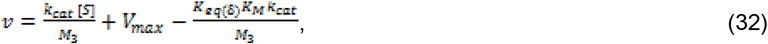

where *k*_*cat*_ *M*_3_, and *v* is the catalytic constant, molar of the substrate, and velocity of product formation respectively. Details leading to Eq. (32) are available in book submitted for evaluation. Taking the intercept as *v*_0_ and the slope (∂*v*/∂[*S*]) being *k*_*cat*_/*M*_3_, one gets:

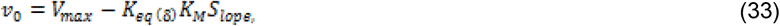

Thus,

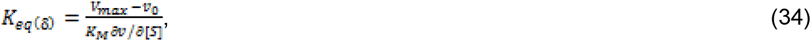

There is also the possibility that the *V*_*max*_ is not known or given but the *K*_*d*_ or *K*_*M*_, [*L*], and [*R*] are known. In this case [*LR*] can be calculated based on Eq. (27). Thus, to address such scenario, Eq. (32) is restated as:

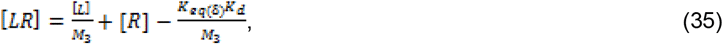

Taking *v*_0L_ as the intercept from the plot of [*LR*] versus [*L*] gives:

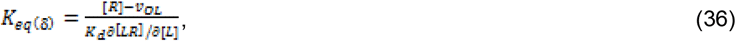

### 2.10. Viral load (*N*_*v*\*(t=i)_) per cell in time (i), later than zero time as a function of the ratio of the experimental viral load (*N*_*v*(τ=0)_) per ml at zero time to the remaining cell/ml after death and the difference of two rate constants-the rate constant (*k*_*v*_) for viral replication and rate constant (*k*_*LR*_) for cell death

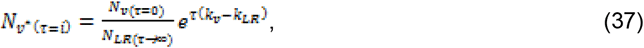

Equation (37) illustrates the relationship between parameters but its simplification requires that *N*_*LR*(τ=0)_*exp*.(−*k*_*LR*_τ) is substituted for *N*_*LR*(τ⟶∞)_ to give:

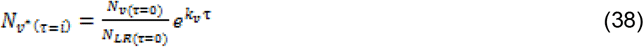

The originating argument leading to Eqs (37) and (38) are available in a book submitted for evaluation.

### 2.11. Viral load (N_v(τ=i)_) per ml as a function of the product of the remaining cell after death and viral load (N_v*(τ=0)_) per cell immediately following infection and the difference of two rate constants (k_v_ and k_LR_)

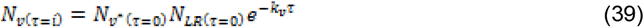

where 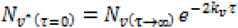

While Eqs (38) and (39) represent the same issue, they are not equal: Their uses in different algebraic setting leads to the same result.

### 2.12. The time (τ_s_) interval between infection and earliest viral load

Given the rate constant for virus infection ((β/copies/mL/cell)/day), viral load at symptom onset (*N*_*v*(τ=0)_/copies/mL), the number of cells (*N*_*LR*_) is given as: First of all, *N*_*v*(τ=o)_ is taken to be directly proportional to *N*_*LR*_ of susceptible host cells. The cells are the kind suggested by Sender *et al*. (2021) in the literature (Crap, *et al*., 1982, Stone, *et al*., 1992), which include bits of information about the number of respiratory system cells, specifically the mucus cells in the nasal cavity (∼10^9^ cells), alveolar macrophages (∼10^10^ cells), and pneumocytes (∼10^11^ cells).

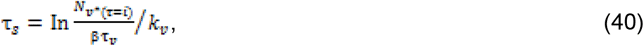

where *N*_*v\** (τ *=* i)_ is defined in Eq. (38) and β is the rate constant for virus infection (Senders *et al*., 2021) as stated earlier, and τ_s_ is the duration between infection and viral load at symptom onset. The alternative equation to Eq. (40) is given as:

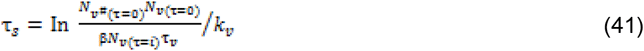

where,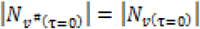, though the unit of the former is viral copies per ml while the latter is viral copies/cell; they may be different but for all practical purpose of evaluating the model, the literature value-the magnitude-of the former is taken to be the same as the latter. Equations (40) and (41) give the same results.

### 2.13. Determination of viral copies/cell/day suitable for different numbers of virus and vulnerable cells with time

As stated earlier, β is used in two ways *viz*: as the same value in viral copies/cell/day and viral copies/ml/day; while stating once again that the two ways may not, in practice, be equal in magnitude the two forms offer a means of evaluating the derived model equations tentatively. Nonetheless, the work of Iwami *et al*. (2012) offers data for the determination of the rate constant for the infection of vulnerable target cells in terms of viral copies/cell/day. The work considers degradation rates of viral RNA, the rate constant for infection of target cells by the virus, viral production rate of infected cells, death rate of infected cells, and even, the growth of target cells. In this investigation, exponential growth due to replication of virus and decrease in the number of cells due to death, are the working models; these therefore, take into account only times when there are increasing trend and decreasing trend in the number of viral RNA and cells respectively. The equation for this purpose is:

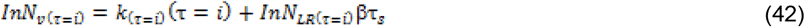

Note that *i* could be zero if the susceptible cell is detectable at such time. If not, *i* > 0. A plot of In *N*_*v*(τ=*i*)_ versus τ=*i*, gives an intercept from which τ_s_ is calculated. The value of β (viral copies/cell/day) is first determined by plotting the initial viral copies per unit time versus the number of cells in time *i*. The slope in the equation given as 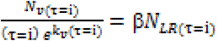 is the value of “β”. Once again all arguments leading to Eq. (42) are available in a book submitted for evaluation.

## 3. EXPERIMENTAL

Frankly speaking, no experiments were conducted. Rather, data in some literature materials were adopted for the evaluation of derived equations. Such citations or references are shown where appropriate tables appear. So, there is no question of materials.

### 3.1. Methods

The methods are completely theoretical, which necessitated the formulation of verifiable equations with which to quantify (or compute) or graphically determine some germane quantities. Despite this, three equations are presented here as substitutes for Eqs. (16) and (22) that yield the same values of *K*_*eq*(δ)_. These substitutes are appropriate for the specified values of [*L*], [*R*], and *K*_*d*_. The concentrations ([*LR*] s) of *LR* complexes were calculated for the spike protein (S-protein)-angiotensin-converting enzyme-2 (ACE2) complex and the receptor-binding domain (RBD)-ACE2 complex. Equation (27), where Σμ_*f*_ (the sum of unequal values of [*L*] and [*R*]) was used in place of 2μ_*f*_, was crucial to this endeavor.

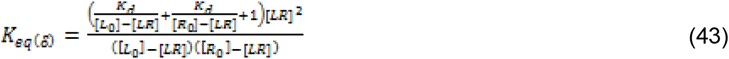

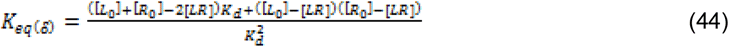

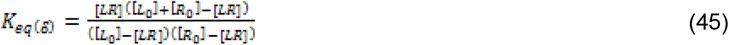

In all equations, Eqs (43), (44), and (45), the subscript 0 refers to the initial concentrations of the species when the time is equal to zero. Moreover, the assumption made is that there are free *R* and *L* at equilibrium thereby calling for the subtraction of [*LR*] from [*R*] and [*L*].

## 4. DISCUSSION

Cholesterol decreases the fluidity of cell membranes and increases their viscosity, but the details of this feature clearly depend on the thermodynamic conditions (Fabian, *et al*., 2023). Cholesterol is known to induce membrane ordering and tighter packing of cell membranes, resulting in slower lateral diffusion (Rog, *et al*., 2009), while at low temperatures, where cholesterol breaks the structure of the gel phase, the situation is just the opposite—a trend in loose packing (Schachter, *et al*., 2022). The presence of proteins and attendant crowding can change the membrane structure (lipid packing and the membrane morphology) within the hydrophobic core and at the polar interface (Tabaka, *et al*., 2014); this causes higher viscosity of the membrane in which the microdomains exhibits much less fluidity than any other part of the membrane. In some cases, the level of lipids is higher than that of different proteins, yet the mobility of proteins within the lipid bilayer is limited to two-dimensional diffusion and is largely determined by the bilayer viscosity (Spei, *et al*., 2011). Sorting and clustering of membrane proteins in a concentration-dependent manner due to lipid dimensions and physico-chemical properties (Marenduzzo, *et al*., 2006, Mazza, *et al*., 2012) can increase the viscosity in the membrane (Tabaka, *et al*., 2014). Membrane viscosity is modulated by the membrane composition, and it is a measure for fluidity (Faizi, *et al*., 2022). This is why lipid raft is very important in the binding of viruses to membranes. The bottom line is that higher viscosity of the lipid rafts enhances the binding of the sSARS-CoV-2.

It is hypothesized therefore, that among any other antidotes, it may be reasonable to deny the invading SARS-CoV-2 a stable base that enables it to inject or release viral material into the cell if and only if the cholesterol and possibly protein content of the lipid raft, estimated to be 20 protein molecules per raft (Simons and Ehehalt, 2002), is reduced. In this regard, studies suggest that increasing cholesterol efflux capacity of HDL could potentially reduce the severity of COVID-19 by affecting the translocation of the ACE2 receptor (including cholesterol-depletion of lipid raft) to lipid rafts and hence the ability of the virus to enter cells (Stadler, *et al*., 2022).

Furthermore, the observed removal of plasma membrane cholesterol following treatment with methyl-β-cyclodextrin, caused the reduction in the levels of ACE2 and the furin protease in lipid rafts, thereby reducing SARS-CoV-2 infection; on the other hand, loading cells with cholesterol via treatment with apolipoprotein (Apo) E and serum increased the trafficking of ACE2 and the furin protease to lipid rafts, resulting in increased SARS-CoV-2 infection (Wang, *et al*., 2021). The binding of sSpike to ACE2 triggers conformational changes that promote the fusion between the host cell membrane and the virus envelope, thereby allowing the viral RNA to be released into the host cell. Compared to other coronaviruses, the spike proteins from SARS-CoV-2 have a very low dissociation constant (14.7 nM) for the binding to ACE2, which makes SARS-CoV-2 highly infectious (Lan, *et al*., 2020).

The irony, based on the experimental outcome, is that the presence of proteins, the spike proteins (sSpike), induces the removal of membrane lipids, both in the presence and absence of ACE2, suggesting that sSpike molecules strongly associate with lipids and strip them away from the bilayer *via* a non-specific interaction. The obvious implication is that it is not thermodynamically feasible for the same cells to become infected twice. The presence of the protein that produces a remarkable degradation of the lipid bilayer (Luchini, *et al*., 2021) can culminate in cell death. In this regard, there is the possibility of inflammatory cell death due to disassembly of AJs and TJs in both AECs and ECs, and hyperplasia following infection. This is in addition to ECM remodeling and deposition of fibrin clots in the alveolar capillaries, leading to disintegration and thickening of the BGB, and ultimately, hypoxia (Makanya, *et al*., 2013).

A cooperative effect of sACE2 and sSpike on lipid extraction has also been observed (Luchini, *et al*., 2021). Scavenger receptor B type I (SR-BI), primarily known as the receptor for HDL, enables the selective uptake of cholesterol esters from HDL particles, thereby contributing to the regulation of cellular cholesterol homeostasis. The role of HDL could be the deposition of cholesterol in the cytoplasm while its efflux role may deplete cholesterol. Intriguingly, SR-BI also facilitates the ACE2-dependent entry of SARS-CoV-2 (Wei, *et al*., 2020) into the cells.

HDLs act as a bridge between the virus and SR-BI since SARS-CoV-2 cannot bind to SR-BI directly (Rani, *et al*., 2024), albeit a contrary view by Alkazim *et al*. (2023) that SARS-CoV-2 binds SR-B1 directly. The binding of the S1 subunit of SARS-CoV-2 to cholesterol and other HDL components, such as apolipoprotein D (Lan, *et al*., 2020), occurs before fusion. There is the possibility that decreasing HDL particle concentrations could serve as a therapeutic strategy to minimize SARS-CoV-2 infectivity (Rani, *et al*., 2024).

The functions and composition of heterogeneous lipoproteins (HDLs) alter in response to changes in an individual’s physiological and/or pathological conditions. This can be described as “chameleon-like characteristics” detrimental to the health of the individual even if the main roles of HDL are cholesterol efflux (including efflux from lipid rafts), anti-inflammatory, and antioxidant functions but become a conveyor of pro-inflammatory agents following its change in composition. This is notwithstanding the fact that molecular intermediates that influence inflammatory microenvironments and cell signaling pathways modulate HDL structural modification and function (Grao-Cruces, *et al*., 2020).

### 4.1 SARS-CoV-2 receptors and entry mechanisms of SARS-CoV-2 by means of its spike protein

Apart from lipids, proteins play a strong role in cellular physicochemical properties, such as the viscosity of the membranes. Some of the proteins are regular members of the lipid raft, while others are not regular members. The fluidity of the membrane is attenuated by the presence of the proteins, some of which are either surface (peripheral protein such as ACE-2) proteins or integral or transmembrane proteins such as Syndecans, a protein of the proteoglycan family (Gopal, 2020), that facilitate the SARS-CoV-2 entry into cells (15). Other proteins, caveolins or clathrins located in the lipid-rich microdomains, the lipid rafts, mediate the entry of enveloped and non-enveloped viruses into host cells, which occurs through fusion or endocytosis (Chazal and Gerlier, 2003). Fusion is often termed early entry mode, while the endocytosis is termed late entry mode. It seems that, regardless of the mode of entry, there must be initial interaction between the viral spike protein and a receptor. Viral infection-binding and injection, viral RNA replication, and cell death is illustrated in a graphic picture (Figure 2) by Palacios-Rápalo, *et al*., (2021) which summarizes all the processes leading to ultimate death of cells.

Let’s consider a moment when there is a temporary equilibrium between the cause of viral infection and the emergence of symptoms. The receptor (*R*) is usually the ACE-2 domicile in the lipid raft. In the first place, Figure 2 by Palacios-Rápalo *et al*. (2021) presents a summary of events beginning from viral-host cell membrane contact and interaction, entry into the by two modes, replication of viral RNA till death of cells. The invading pathogen is the virus spike protein regarded as the ligand (*L*). It is to serve as pictorial background to the relevance of all models (equations) for the quantification of thermodynamic parameters and for the use of available literature kinetic parameters to generate dimensionless equilibrium constant. In binding processes, the surface-exposed S1 part of the homotrimeric class I fusion glycoprotein containing the receptor-binding domain (RBD) interacts with host cell receptor, thereby determining virus cell tropism (the capacity of a virus to infect its target) and pathogenicity. The second part, the transmembrane S2 domain contains heptad repeat regions and the fusion peptide (Wei, *et al*., 2020, Clausen, *et al*., 2020, Wang, *et. al*., 2020). Infection begins strongly when, after invasion, the pathogens are able to bind to the host cell membranes. Binding yields a receptor-ligand complex.

The spike protein consists of two subunits—the S1, which interacts with ACE-2, and the S2, which mediates the fusion. The furin cleavage motif that lies between S1 and S2 is cleaved by the host cell’s transmembrane serine protease (TMPRESS2) at the S2 site following the binding of the virus to the membrane. The cleavage facilitates the fusion of the S2 subunit with the host cell bilayer, releasing the virus RNA into the cells. Nonetheless, the concentration of TMPRESS2 may be too low if expressed at all, and in such a situation, the virus explores the endosomal pathway using ACE-2 and cathepsin. Once the viral RNA is in the cell, the host protein production machinery is hijacked through what has been described as a cross-talk with autophagic machinery and mitochondrial metabolism (Palacios-Rápalo *et al*. 2021). The term cross-talk may be defined as the effect or influence of one or more components of a signal transduction partway on other. This also refers to the control of a subcellular system by another biotic (genetic and microbial entities) or xenobiotic system. Such influence could be unbeneficial by being harmful; it may be beneficial if the pathogen is undermined by the xenobiotic.

It is likely that each cell membrane contains a number of lipid rafts that contain *ACE*2, which suggests that many viruses attach to the host cell membrane. It has been hypothesized that for a raft of size 50 nm, the number of lipid rafts is equal to 10^6^ (Simons and Ehehalt, 2002). It is quite obvious that binding of any virus to membranes is to specific domains containing specific receptors; low *K*_d_ suggests strong affinity of spike protein to its receptor, ACE-2, and other receptors. Thus, Eqs (17) and (20) for free energy of binding and activation energy, respectively, can quantify the corresponding parameter given pieces of information about the concentrations of spike protein, ACE-2, and other receptors, as well as spike protein-ACE-2 complex. Iwani *et al*. (2012) identified events such as the degradation-rate of viral RNA, rate constant for virus infection of host target cells, death rate of infected cells (1.75/day)

While admitting that Chol on cell membranes facilitates viral entry, the role of intracellular Chol in SARS-CoV-2 cannot be overemphasized: Genes (from CRISPR libraries) including sterol-regulatory element-binding protein (SREBP-2), SREBP cleavage-activating protein (SCAP), low-density lipoprotein receptor (LDLR), and membrane-bound transcription factor peptidases, site 1 and 2 (MBTPS 1 and MBTP 32), for cholesterol metabolism are essential for SARS-CoV-2 infection. Besides, treatment with amlodipine, a calcium ion channel antagonist, increases intracellular cholesterol levels, thereby significantly inhibiting SARS-CoV-2 infection (Kluck *et al*., 2021).

### 4.2. Dimensionless equilibrium constant for the binding of viral particle, the ligand to the receptor, the ACE2

In other to determine the dimensionless equilibrium constant for the binding of viral particle, the ligand to the receptor, the ACE2, the values of [*LR*] were theoretically determined using Eq. (27). The values of [*L*], the concentration of spike protein is given in the literature [68] and explored in this research. Included in the same literature is the equilibrium dissociation constant, *K*_*d*_. Of course, the *LR* complex represents the spike protein-ACE2 complex for the wide type (WT) and D614G variants. The [*LR*] were displaced in Table 1.

The graphical determination of the dimensionless equilibrium constant, *K*_*eq* (δ),_ based on Eq. (35), is by plotting [*LR*] versus [*L*] on the assumption that the receptor, *R*, has a constant mass concentration equal to 200 ng/ml (Ozono, *et al*., 2021). The *K*_*eq* (δ)_ values were calculated using Eq. (36). The plots were for temperatures of 298.15, 303.15, and 310.15 K for the wild type and 614G variant. Figures 3, 4, and 5 were for the wild type, while Figures 6, 7, and 8 were for the 614G variant. The plots exhibited a hyperbolic curve characteristic of Michaelian kinetics. This is shown as inset in all the graphical figures covering the temperature range. Therefore, the linear phase covering the lower concentrations of the ligand was adopted for each plot; the implication is that the equations stated earlier are suitable for the conditions that go with the reverse quasi-steady-state approximation (rQSSA) and total QSSA (tQSSA), referring to a scenario where [*R*] > [*L*] or [*R*] ≈ [*L*].

**Figure 3:**
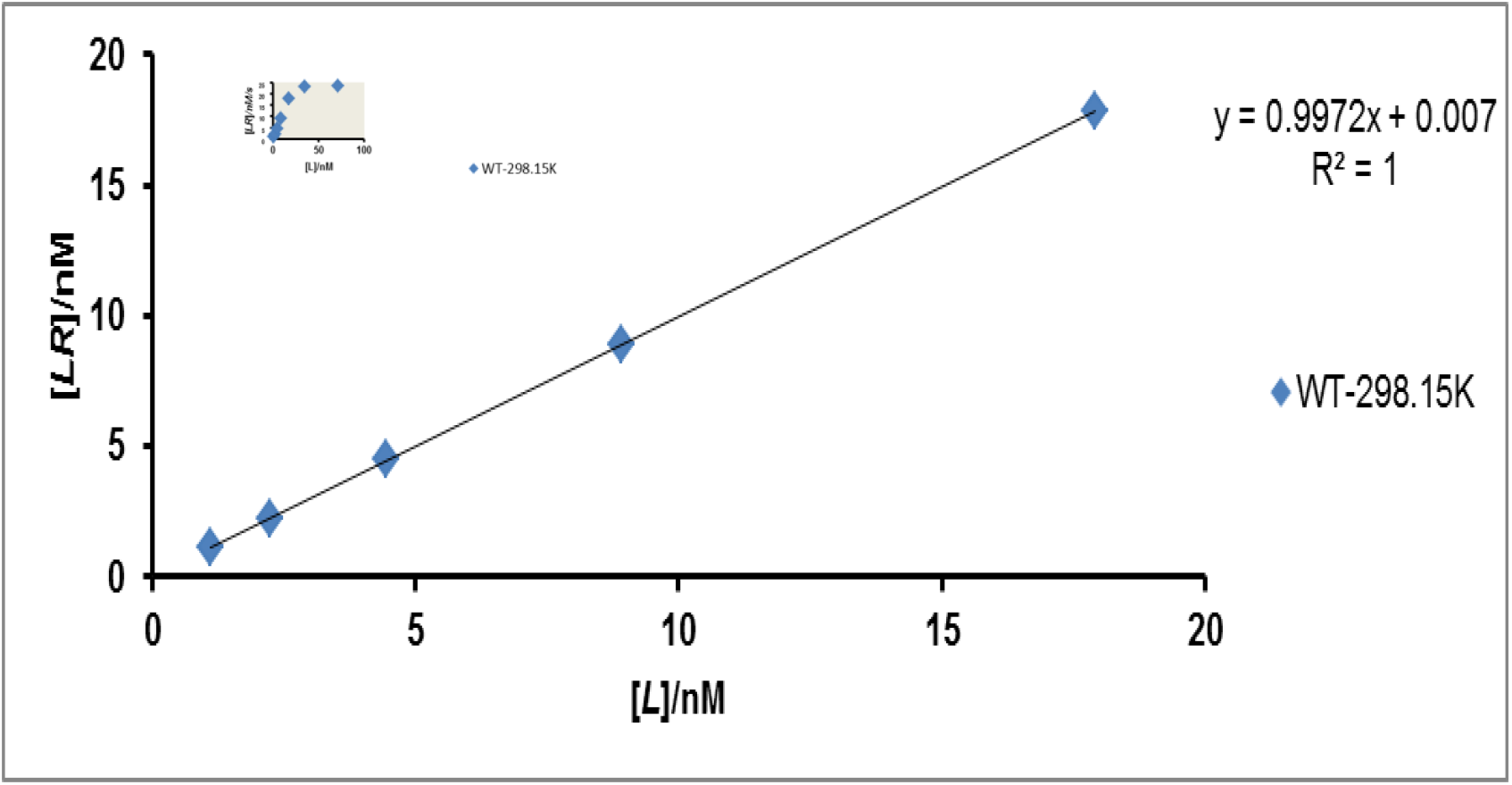
Determination of dimensionless equilibrium constant based on Eqs (35) and (36) for wild type (WT) at 298.15 K. The molar concentration of ACE2 is equal to 23.28140509 nM on the assumption that the mass per unit volume was 200 ng (Ozono, *et al*., 2021); the molar mass of ACE2 is equal to 85.9 kDa. Any other data are from the same reference.

**Figure 4:**
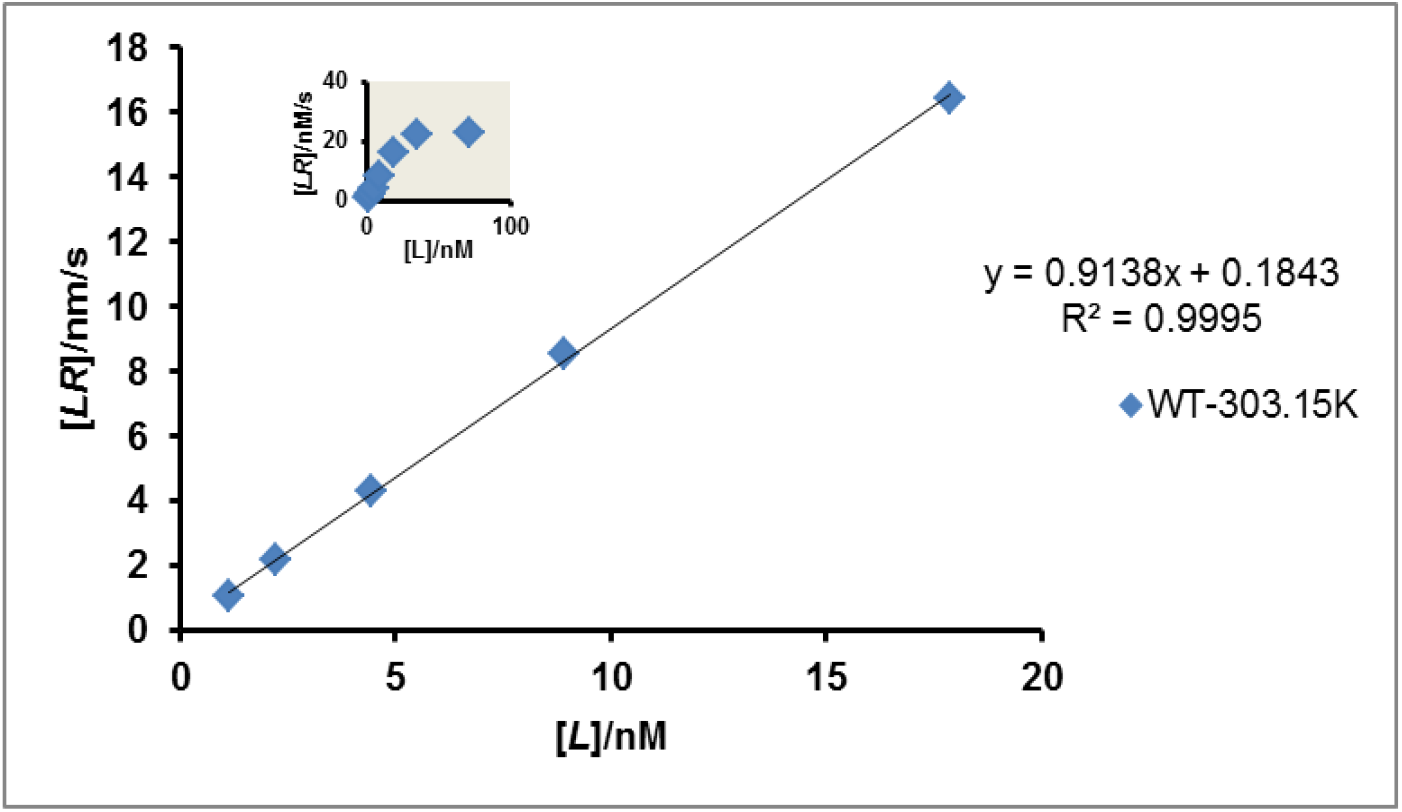
Determination of dimensionless equilibrium constant based on Eqs (35) and (36) for wild type (WT) at 303.15 K. The molar concentration of ACE2 is equal to 23.28140509 nM on the assumption that the mass per unit volume was 200 ng (Ozono, *et al*., 2021); the molar mass of ACE2 is equal to 85.9 kDa. Any other are from the same reference

**Figure 5:**
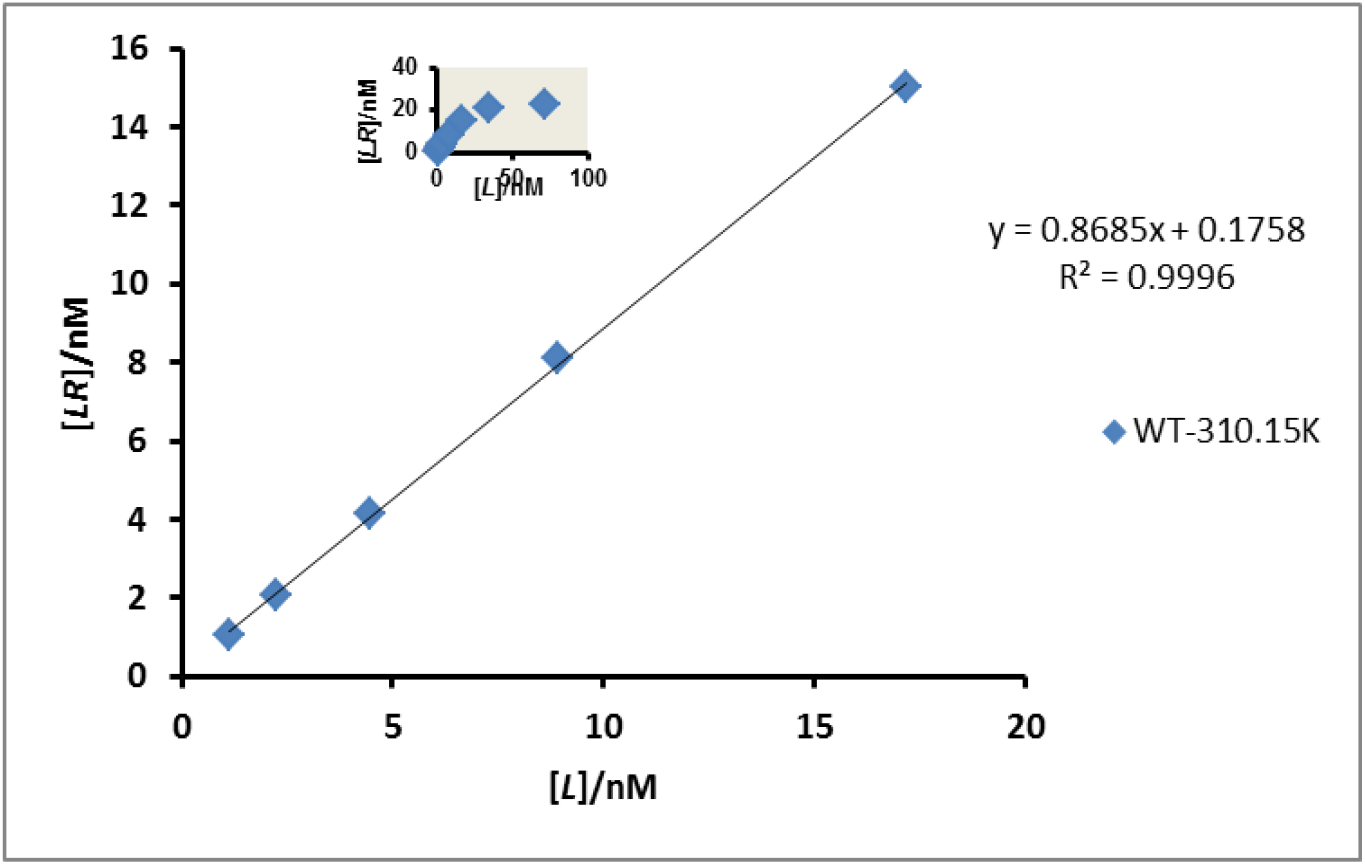
Determination of dimensionless equilibrium constant based on Eqs (35) and (36) for wild type (WT) at 310.15 K. The molar concentration of ACE2 is equal to 23.28140509 nM on the assumption that the mass per unit volume was 200 ng (Ozono, *et al*., 2021); the molar mass of ACE2 is equal to 85.9 kDa. Any other are from the same reference

**Figure 6:**
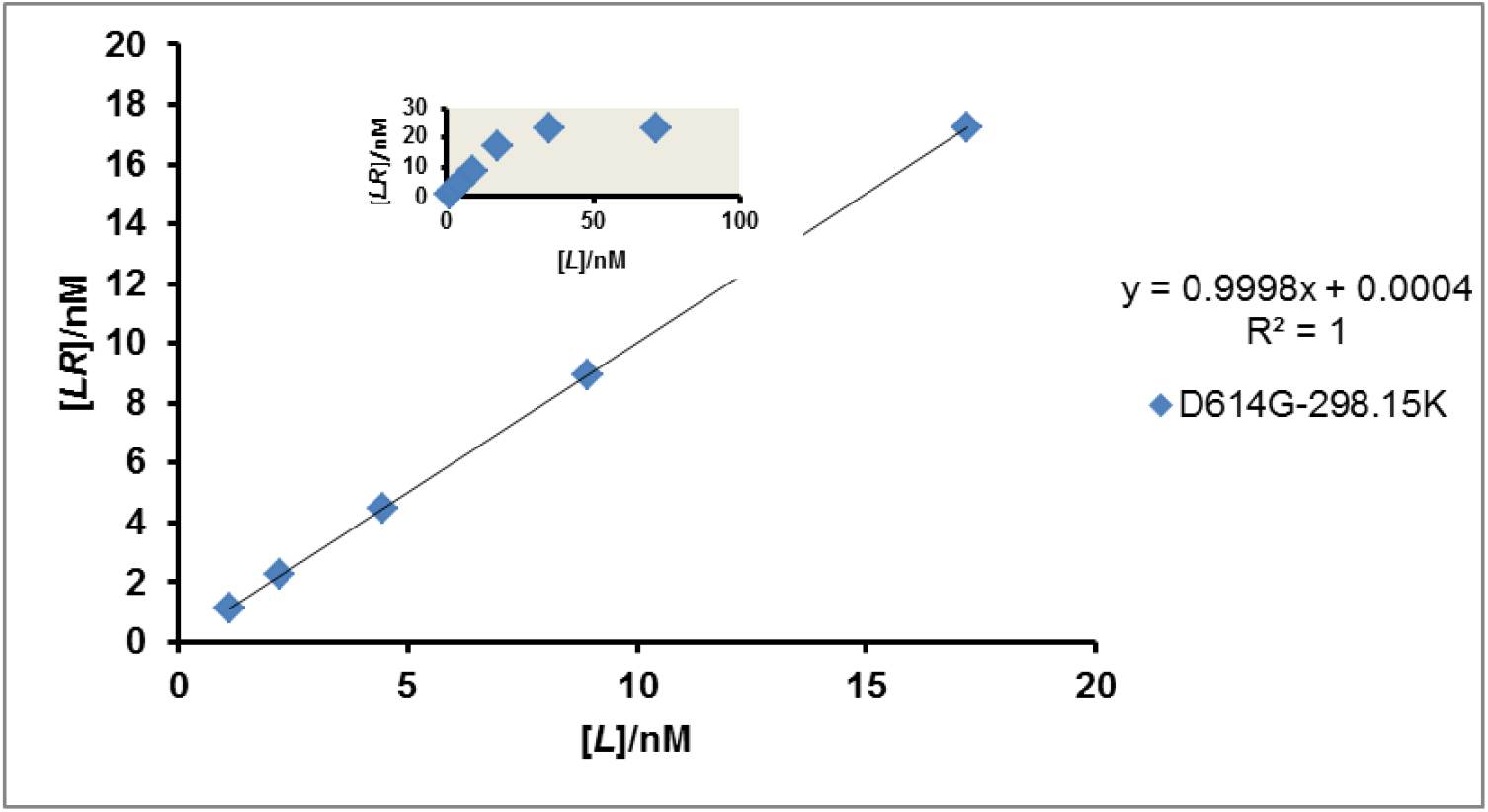
Determination of dimensionless equilibrium constant based on Eqs (35) and (36) for D614G at 298.15 K. The molar concentration of ACE2 is equal to 23.28140509 nM on the assumption that the mass per unit volume was 200 ng (Ozono, *et al*., 2021); the molar mass of ACE2 is equal to 85.9 kDa. Any other data are from the same reference.

**Figure 7:**
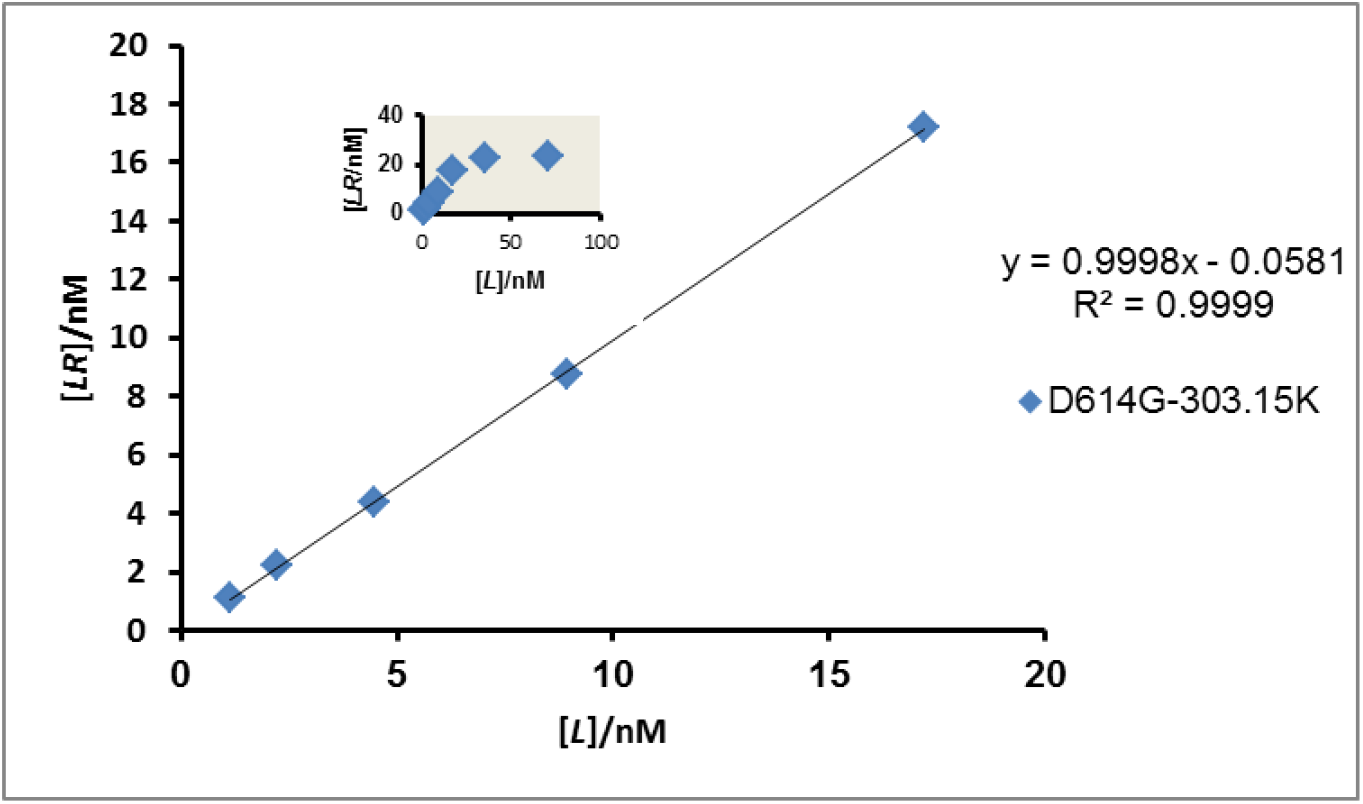
Determination of dimensionless equilibrium constant based on Eqs (35) and (36) for D614G at 303.15 K. The molar concentration of ACE2 is equal to 23.28140509 nM on the assumption that the mass per unit volume was 200 ng (Ozono, *et al*., 2021); the molar mass of ACE2 is equal to 85.9 kDa. Any other data are from the same reference.

**Figure 8:**
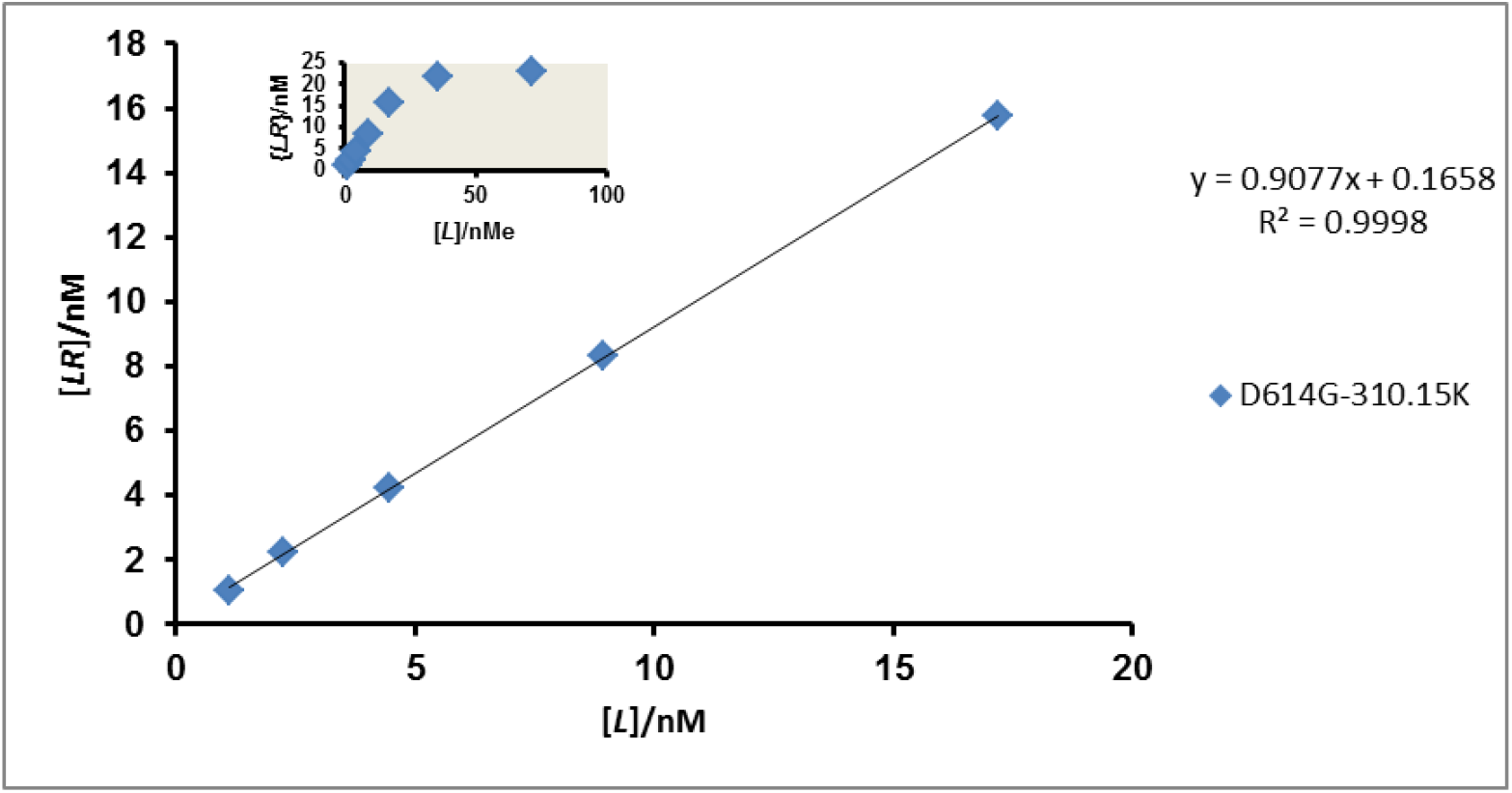
Determination of dimensionless equilibrium constant based on Eqs (35) and (36) for D614G at 310.15 K. The molar concentration of ACE2 is equal to 23.28140509 nM on the assumption that the mass per unit volume was 200 ng (Ozono, *et al*., 2021); the molar mass of ACE2 is equal to 85.9 kDa. Any other data are from the same reference

In order to compute the dimensionless equilibrium constant, *K*_eq(δ)_, Eq. (36) was explored by fitting it to the slopes and intercepts from the plots, Figures 3 through 8. The equation is also fitted to experimentally generated dissociation constant (*K*_*d*_) values such as (Ozono, *et al*., 2021): 1.46 *e*. (−11) M at 298.15 K; 6.14 *e*. (−10) M at 303.15 K; 1.58 *e*. (−9) M at 310.15 K for the WT. Other values are: < 1 *e*. (−12) M at 298.15 K; 2.97 e. (−10) M at 303.15 K; 1.04 *e*. (−9) M at 310.15 K for the D614G. The results of such computation are displayed in Table 2. As could be seen, the *K*_eq(δ)_ values (Table 2) decrease with increasing thermodynamic temperature. The values for the D614G variant are higher than WT values; the corresponding Gibbs free energies of spike protein-ACE2 complex formation are also higher in negative magnitude than values for WT variants. The thermodynamic feasibility is greater for D614G than for the WT variant, and greater feasibility is more pronounced at lower temperatures than at higher temperatures.

Tables 2 and 3 demonstrate that the patterns of the thermodynamic parameter values with temperature are different; Table 3 breaks the increasing trend in the negative magnitude at 303.15 K, in contrast to Table 2. This can be due to unknown elements in the literature’s methodology (Sepsey, *et al*., 2020, Carrero, 2024). In general, both WT and D614G variants exhibit exothermically (exogonically) or rather, thermally induced spike protein binding to ACE2. A reasonable inference is that ligand-receptor complex formation is very stable. This is greater for D614G than for the WR variant. The enthalpy change was generated from van’t Hoff’s plot (Figure 9).

**Table 3:** Exploring van’t Hoff equation given as: In*K*_*eq*(δ)_ = −Δ*H/RT* + *C* where *C* may be Δ*S/R* (Sepsey, et al., 2020, Carrero, 2024)

|  | WT |  | D614G |  |
| --- | --- | --- | --- | --- |
| **T/K** | **T$\Delta S$/e. (5)** | **$\Delta G$** | **T$\Delta S$/e. (5)** | **$\Delta G$** |
| **298.15** | -2.64 | -1.67 e. (4) | - 4.00 | - 2.24 e. (4) |
| **303.15** | -2.19 | -1.23 e. (4) | - 4.06 | -1.56 e. (4) |
| **310.15** | -2.75 | - 6.06 e. (3) | - 4.16 | -6.24 e. (3) |

WT: $-\Delta H/R = 33797$ , $\Delta S/R = -106.62$ J/K/mol., $\Delta H = -2.81$ e. (5) J/mol., $\Delta S = -886.4$ e. (5) J/K/mol. and D614G: $-\Delta H/R = 50762$ , $\Delta S/R = -161.25$ J/K, $\Delta H = - 4.22$ e. (5) J/mol., $\Delta S = - 1340.68$ e. (5) J/K. Results were approximations to 2 decimal places. Data in the literature (Ozono, *et al.*, 2021) were explored for the generation of the data in this research.

**Table 4:**
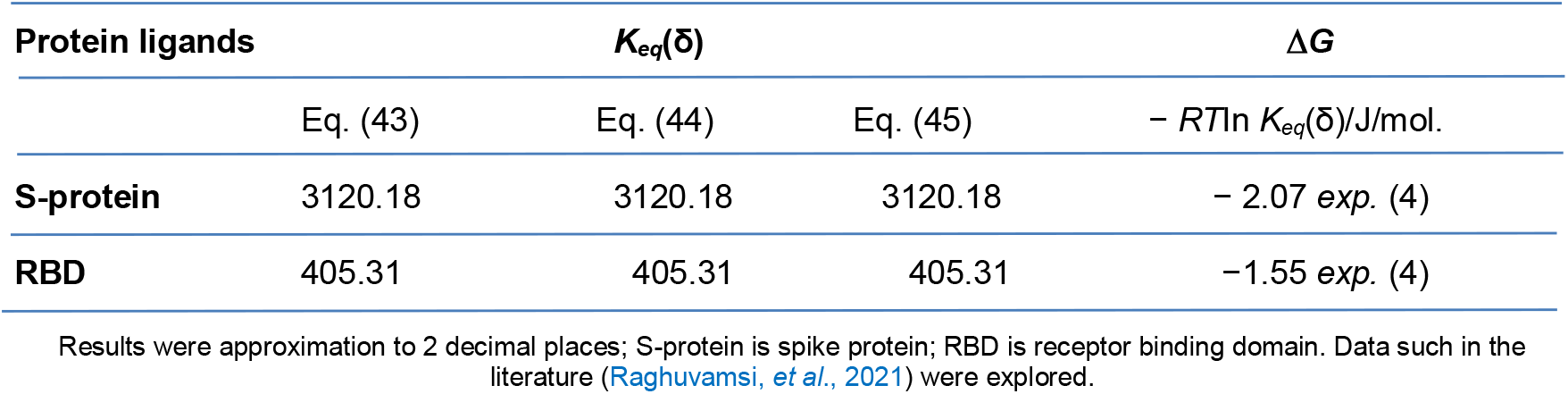
The dimensionless equilibrium constant of binding interaction of the ligands (S-protein and RBD) and the receptor, ACE2 and free energy of binding.

| Protein ligands | $K_{eq}(\delta)$ | | | $\Delta G$ |
| --- | --- | --- | --- | --- |
| | Eq. (43) | Eq. (44) | Eq. (45) | $- RT \ln K_{eq}(\delta)/J/mol.$ |
| **S-protein** | 3120.18 | 3120.18 | 3120.18 | $- 2.07 \text{ exp. (4)}$ |
| **RBD** | 405.31 | 405.31 | 405.31 | $-1.55 \text{ exp. (4)}$ |

Results were approximation to 2 decimal places; S-protein is spike protein; RBD is receptor binding domain. Data such in the literature (Raghuvamsi, *et al.*, 2021) were explored.

**Figure 9:**
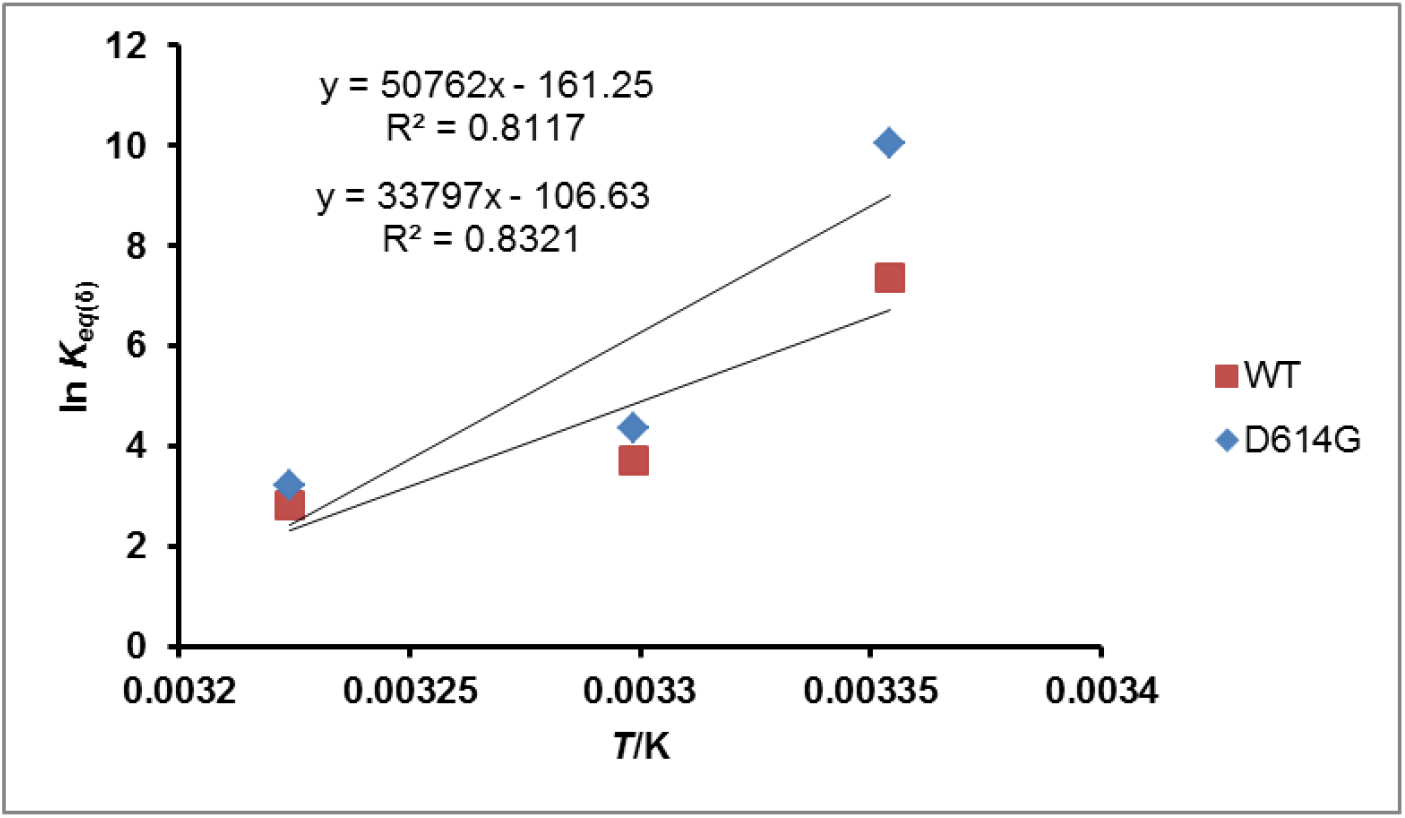
Determination of enthalpy of binding of S-protein to ACE2 for the wide type WT (Brown Square) and D614G (Blue rhombus). Δ*H*^0^: 4.22050497 *e*. (5) J/mol. (D614G); 2.80998397 *e*. (5) J/mol. (WT). The original data explored are as in the literature (Ozono, *et al*., 2021).

Next, attention should be shifted to activation energy issues. Briefly, let us recall the meaning of activation energies referred by Copeland (page 153) (Copeland, 2002) in the works of Ferst *et al*. (1974) and So, *et al*. (1988). The overall activation energy *E*_a_ is composed of two terms, Δ*G*_*ES*_ and 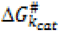. The term 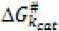 is the amount of energy that must be expended to reach the transition state (*i*.*e*., bond-making and bond-breaking steps), while the term Δ*G*_*ES*_ is the net energy gain that results from the realization of enzyme*—*substrate (or ligand*—*receptor) binding energy (Ferst, *et al*. 1974, So, *et al*. 1988). In this research, the focus is on the Gibbs free energy of activation for the dissociation 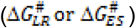 of *LR* or *ES* complex in reverse order. However, the Arrhenius equation is explored for the determination of activation energy as shown in Figure 10. The reverse first-order dissociation constant (*k*_*dis*_) values corresponding to different temperatures (298.15*—*310.15 K) for the *LR* complex in the literature (Ozono, *et al*., 2021) were explored. It was however, observed that the *k*_*dis*_ values for WT (298.15 K: 1.82 *e*. (−6)/s; 303.15 K: 97 *e*. (−6)/s; 310.15 K: 300 *e*. (−6)/s) and D614G (298.15 K: < 0.1 *e*. (−6)/s; 303.15 K: 53.9 *e*. (−6)/s; 310.15 K: 168 *e*. (−6)/s) protein variants as ligands were increasing with increasing temperature with the values for D614G being lower than the values for WT.

**Figure 10:**
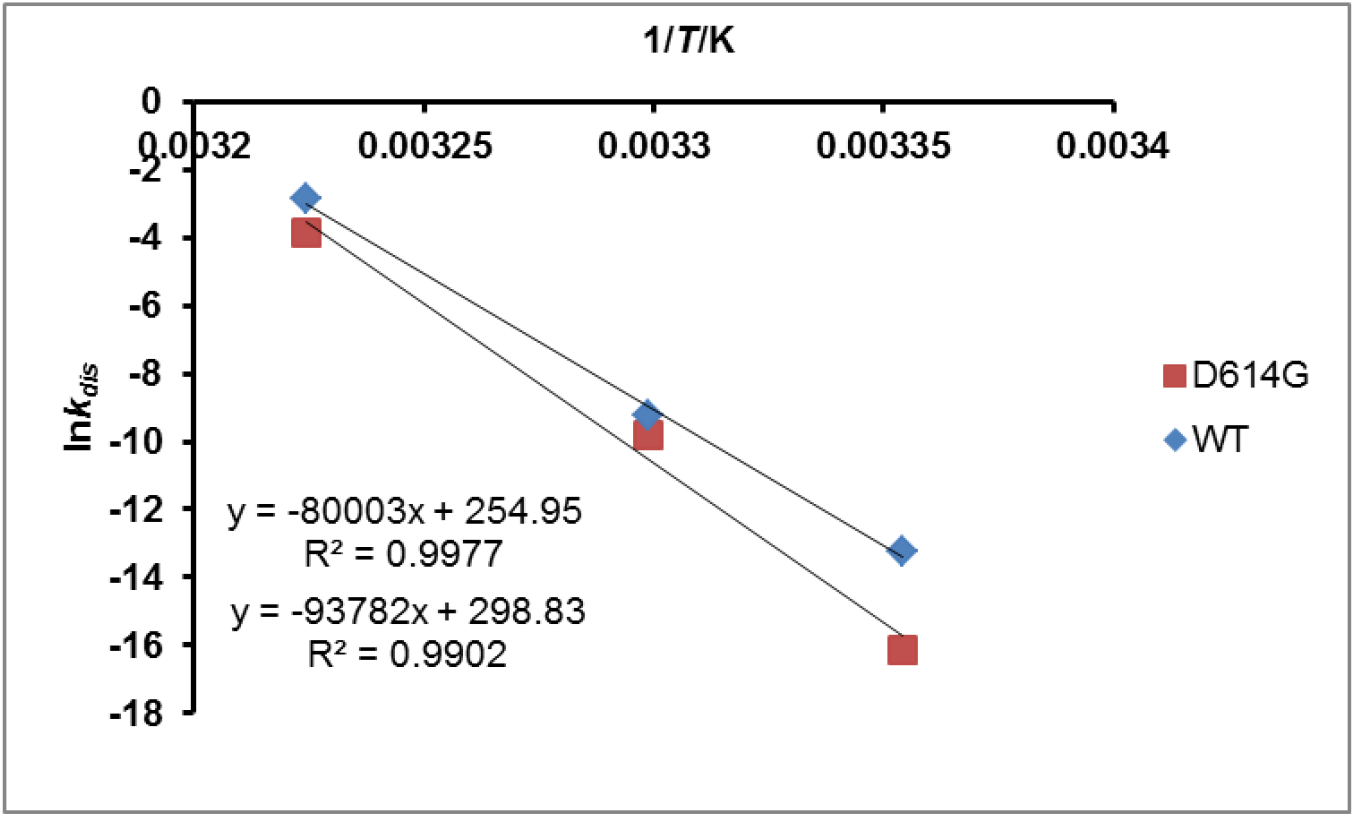
Activation energy (Arrhenius type) for the dissociation of S-protein-ACE2 complex. The square in brown color stand for D614G while the rhombus stands for WT. WT: 6.65 *e*. (5) J/mol. and D614G: 7.89 *e*. (5) J/mol. Both are approximations to 2 decimal places. The original data explored are as in the literature (Ozono, *et al*., 2021)

The implication is that the 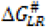 values could be lower at higher temperatures; this therefore, suggest that the amount of energy to be expended to accomplish the transition state before dissociation of the *LR* complex in reverse order is lower at higher temperatures with the result that the *LR* is less stable at higher temperatures than at lower temperatures. The overall implication is that the *LR* complex for the WT protein variant is less stable than *LR* complex for D614G. This view is in line with the conclusion that the D614G mutation increases cell entry by acquiring higher affinity than the WT to ACE2 (Ozono, *et al*., 2021). It should not be a surprise that by the same argument, the Arrhenius type of activation for the dissociation of the *LR* complex (Figure 10) for D614G is higher than for WT (WT: 6.65 *e*. (5) J/mol. and D614G: 7.89 *e*. (5) J/mol.).

There is no doubt that the issue of binding free energy has taken a front burner in the scheme of things connected with viral infection. The reason is that the spontaneity of binding of the ligand to receptor expresses its thermodynamic feasibility that is often signaled by the magnitude of the equilibrium dissociation constant, *K*_*d*_. Examples of such *K*_*d*_ values are available in the literature (Nguyen, 2020, Ozono, *et al*., 2021). With such data it has been possible for comparison between two variants of coronavirus to be made. For instance, it has been observed that the RBD of SARS-CoV (SARS-CoV-RBD) has lower affinity for human ACE2-PD than SARS-CoV-2-RBD. Although different groups may report different values of *K*_*d*_ for the same variants studied (Nguyen, 2020), there are bits of evidence for the differences between variants. For instance, for SARS-CoV, Kirchdoerfer, *et al*. (2018) and Wrapp, *et al*. (2019) reported *K*_*d*_ ≈ 150−300 nM, which is 10−20 times greater than that of SARS-CoV-2. This implies that the old virus (SARS-CoV) is much more weakly associated with ACE2 than the new virus (SARS-CoV-2). Whereas, Walls, *et al*. (2020) is of the view that *K*_*d*_ of SARS-CoV is about 4 times larger than *K*_*d*_ of SARS-CoV-2, while a simulation by Nguyen, *et al*. (2020) yielded an approximately 2 times larger value for SARS-CoV than for SARS-CoV-2.

There is a need to opine too that unlike the very frequent reference to osmophobic effect of osmolyte (the unfavourable interaction of the organic osmolyte with peptide back bone) which must have a basis (often not specified to explain exclusion of the osmolyte), the binding of spike protein or any other protein ligand to the receptor is driven by electrostatic interaction which should be specifically attractive (Nguyen, 2020). The binding of spike protein or any other protein ligand to the receptor is driven by electrostatic interaction, in contrast to the often mentioned osmophobic effect of osmolyte (the unfavorable interaction of the organic osmolyte with peptide backbones), which requires a basis (often not specified to explain the exclusion of the osmolyte). This ought to be very appealing.

If, according to convention, the greatest potential energy between two sites where oppositely charged particles are placed is zero, then the numbers with negative signs appear to indicate that the potential energies were recorded. In addition, the system’s total energy is equal to the kinetic energy of attraction but has the opposite sign. These values, as well as van der Waal values, assuming total energies are: − 620.39 and − 791.41 kcal/mol. for SARS-CoV and SARS-CoV-2 respectively. The van der Waals components follow the same order: − 75.56 and −84.48 kcal/mol. for SARS-CoV and SARS-CoV-2 respectively (Nguyen, *et al*., 2020). This could help to clarify why SARS-CoV-2 spike protein binds to the receptor more firmly than SARS-CoV. In addition to the energetic contributions, scientists (Ali and Vijayan, 2020) are interested in the free energy of binding (Δ*G*_*bind*_) of SARS-CoV-2 and SARS-CoV S RBDs to ACE2. They estimated these contributions from relatively unknown (unfamiliar) frames of all MD simulations using the molecular mechanics-generalized Born surface area (MM-GBSA) approach; the reported values for the binding of SARS-CoV-2 SRBD and SARS-CoV SRBD to ACE2 range between −106 and −140 and −44.5 and −71.2 kcal/mol, respectively. Even though these values are far larger than those calculated for WT and D614G, they nevertheless, provide more proof of variations in binding affinity and its free energy component.

The major issue is that the *K*_*d*_ numbers cannot be utilized directly as Δ*G*_*bind*_ = −*k*_*B*_*T* ln(*K*_*d*_) to compute the free energy of binding since *K*_*d*_ values have units. If the molar concentrations of SARS-CoV-RBD, SARS-CoV-2-RBD, and ACE2-PD are known, the value of the *LR* complex concentration can be estimated as described earlier (Eq. (27)). After that, *K*_*eq*(δ)_ can be calculated by fitting Eq. (46) or any other equation to pertinent data. Comparing thermodynamic parameters based on the techniques used in this study with the findings that are based on *K*_*d*_ values is not appropriate at this time. Nonetheless, the latter’s findings might show patterns that are comparable to those seen when exploring *K*_*eq*(δ)_ values.

There are undoubtedly several ways to calculate *K*_*d*_, but understanding [*LR*] is essential for more computational work. However, such techniques include combining coarse-grained and all-atom steered molecular dynamics (SMD) simulations (Nguyen, 2020), surface Plasmon resonance approach (Wrapp, 2019), bio-layer interferometry (Walls, *et al*., 2020), *etc*., even though they are not familiar to much less developed countries. These techniques might not preclude methods developed in this research and detailed in a book submitted for assessment.

### 4.3. Thermodynamics of the anabolism of the intermediate (3-OH-3-methyl glutaryl-CoA) in the synthesis of cholesterol

In light of the role of cholesterol in the pathogenesis of COVID-19 due to severe acute respiratory syndrome coronavirus 2 (SARS-CoV-2), any part of the pathway in its synthesis is important. Here is presented, based in part on the data in the literature (Bouchard, *et al*., 2021), the dimensionless equilibrium constants. These were determined by exploring Eq. (35) and Eq. (36). The reaction between acetoacetyl CoA and acetyl CoA is catalyzed by HMG-CoA synthase to yield 3-OH-3-methyl glutaryl-CoA (HMG-CoA), which is an intermediate in the synthesis of cholesterol. A mutant CHS (mutant Chediak-Higash syndrome), which is an organism that has a mutation in the lysosomal traffic regulator gene (Bouchard, *et al*., 2021), and the wild type were adopted for the assay in the production of HMG-CoA. The dimensionless equilibrium constants were 1.1560749855 and 3.367234744 for the wild type and F137L cHS, respectively. Accordingly, the free energies are −365.55 J/mol. and −3060.09 J/mol. These results show that with the F137L variant, the reaction catalyzed with HMG-CoA synthase is more feasible and spontaneous than the wild type.

### 4.4. Thermodynamic characterization of the binding of SARS-CoV and receptor binding domain (RBD) to angiotensin converting enzyme2 (ACE2)

The determination of dimensionless equilibrium constant (*K*_*eq*_(δ)) and cognate free energies explored the data in the literature (Raghuvamsi, *et al*., 2021). The dimensionless equilibrium constants were calculated using Eq. (43), Eq. (44), and Eq. (45). All equations were used to demonstrate their robustness and validity.

### 4.5. The thermodynamics of viral replication and concurrent death of susceptible cells

The thermodynamics of viral replication in terms of higher number of copies per day and cell death are herein given attention. The relevant equation is Eq. (50a). However, the velocity of the production of viral RNA copies (the number of copies per day) may not be the same as the rate constant for viral RNA production. This comment is analogous to the velocity of hydrolysis as opposed to the first-order constant for the same action. Different days present different rates of production of viral RNA copies. This is shown in the literature (Iwami *et al*., 2012). However, it is not very certain the authors meant rate constant. Besides, they stated the unit of the rate constant for infection (β) as (RNA/ml · day) ^−1^. The reciprocal of it is RNA copies/ ml. day, even if the value of β was 8.61 *e*. (−11). This implies that the value should be stated as 8.61 *e*. (−11) (RNA/ml · day) ^−1^. This seeming ambiguity is unlike the units of viral production rate and death rate of susceptible cells, which are, respectively, 3.24 *e*. (4) RNA copies/day and 1.75/day. Nonetheless, it is uncertain whether the different susceptible cells possess the same rate of death. In other to address this challenge, graphical approaches were explored for the determination of death rate constants rather than velocities. Since viral particles replicate in a host cell, the death of cells that follows are contemporary events: The fact that the host genetic material is hijacked is synonymous to cell death. Figures 11 and 12 present the plots showing death rate constant (*k*_*LR*_) 1.1609 and 1.1241/day for Nef-Negative HSC-F Cells and Nef-Positive HSC-F Cells respectively. These values are, however, less than 1.75 /day (Iwami *et al*., 2012), which seemed to be arbitrarily chosen (although it was reported as an estimate) for, perhaps, convenience’s sake. This research sees it differently because the cells are different *ab initio*.

**Figure 11:**
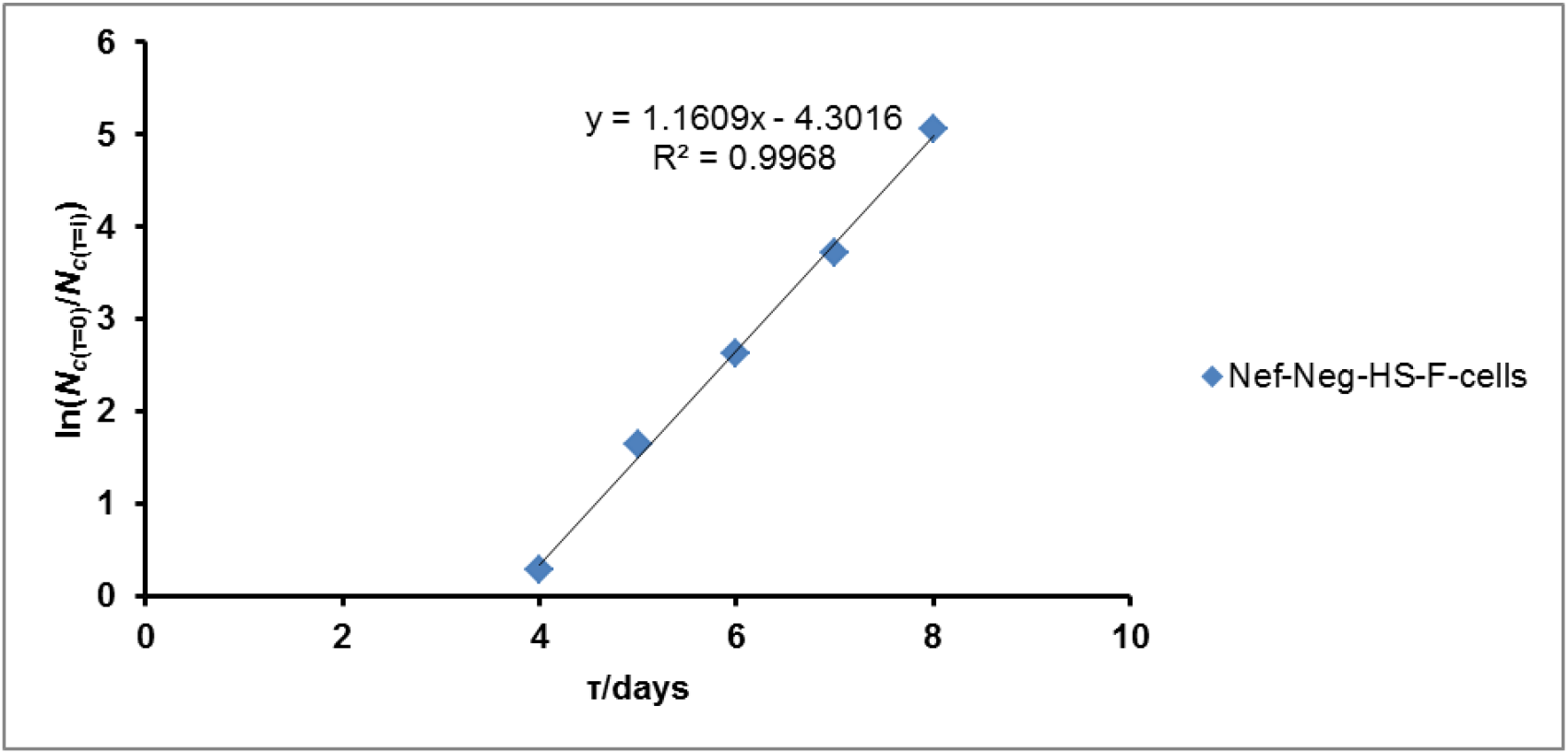
Death rate constant of Nef-Negative HSC-F Cells (Nef-neg-HSC-F-Cells) by a plot of In(*N*_*c*(*τ*=0)_/*N*_*c*(τ=*i*)_) versus the time, τ in days where *i* is any time ≪ ∞. Enabling data are from Iwami *et al*. (Iwami *et al*., 2012).

**Figure 12:**
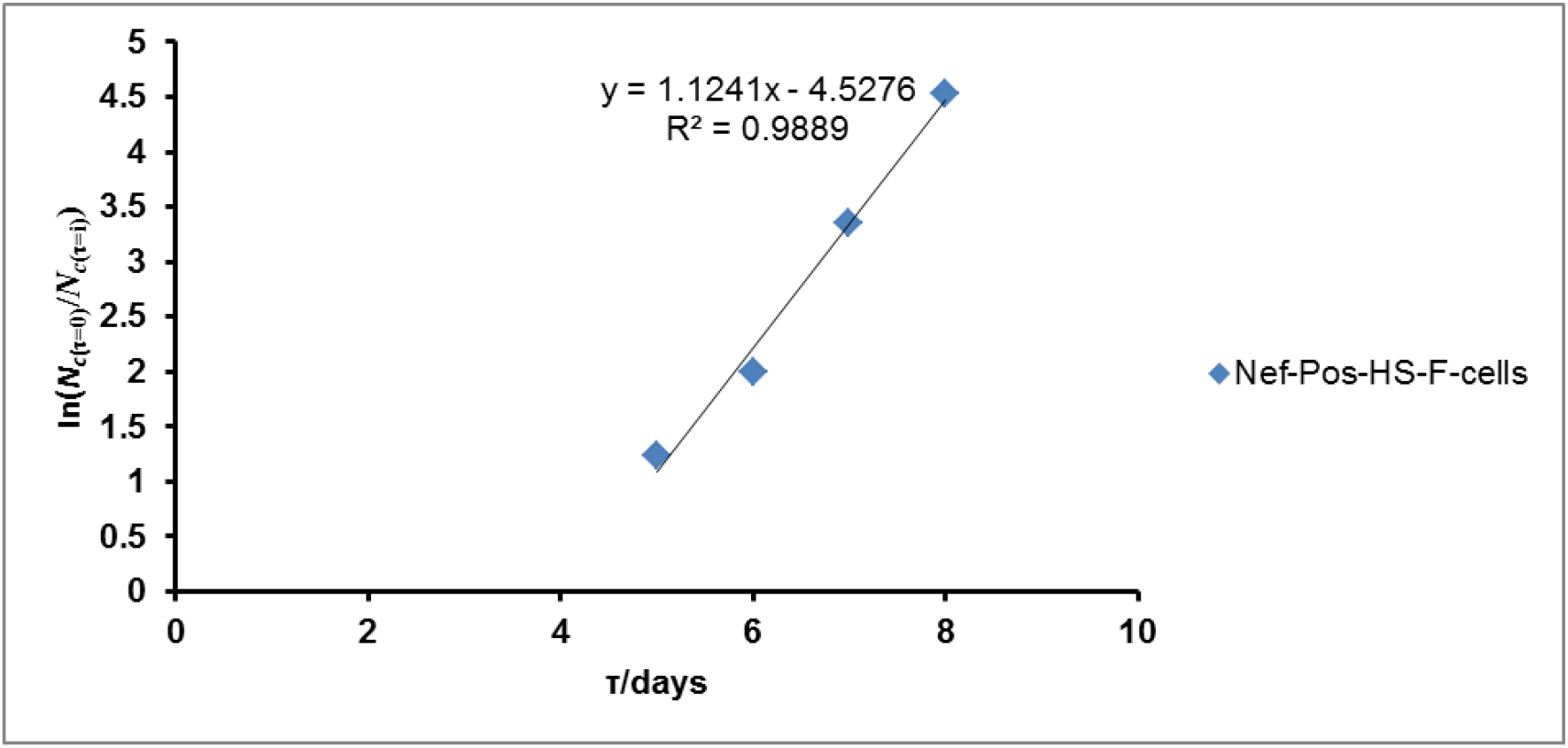
Death rate constant of Nef-Positive HSC-F Cells (Nef-Pos-HSC-F-cells) by a plot of In(*N*_*c*(τ=0)_/*N*_*c*(τ=*i*)_) versus the time, τ in days where *i* is any time ≪ ∞. Reference is as stated earlier.

There was a need to also determine graphically the rate constant for viral replication. Figures 13 and 14 show that, the rate constants for the replication of the virus are 2.5192 and 2.1957/day for Nef-Negative HSC-F Cells and Nef-Positive HSC-F Cells respectively.

**Figure 13:**
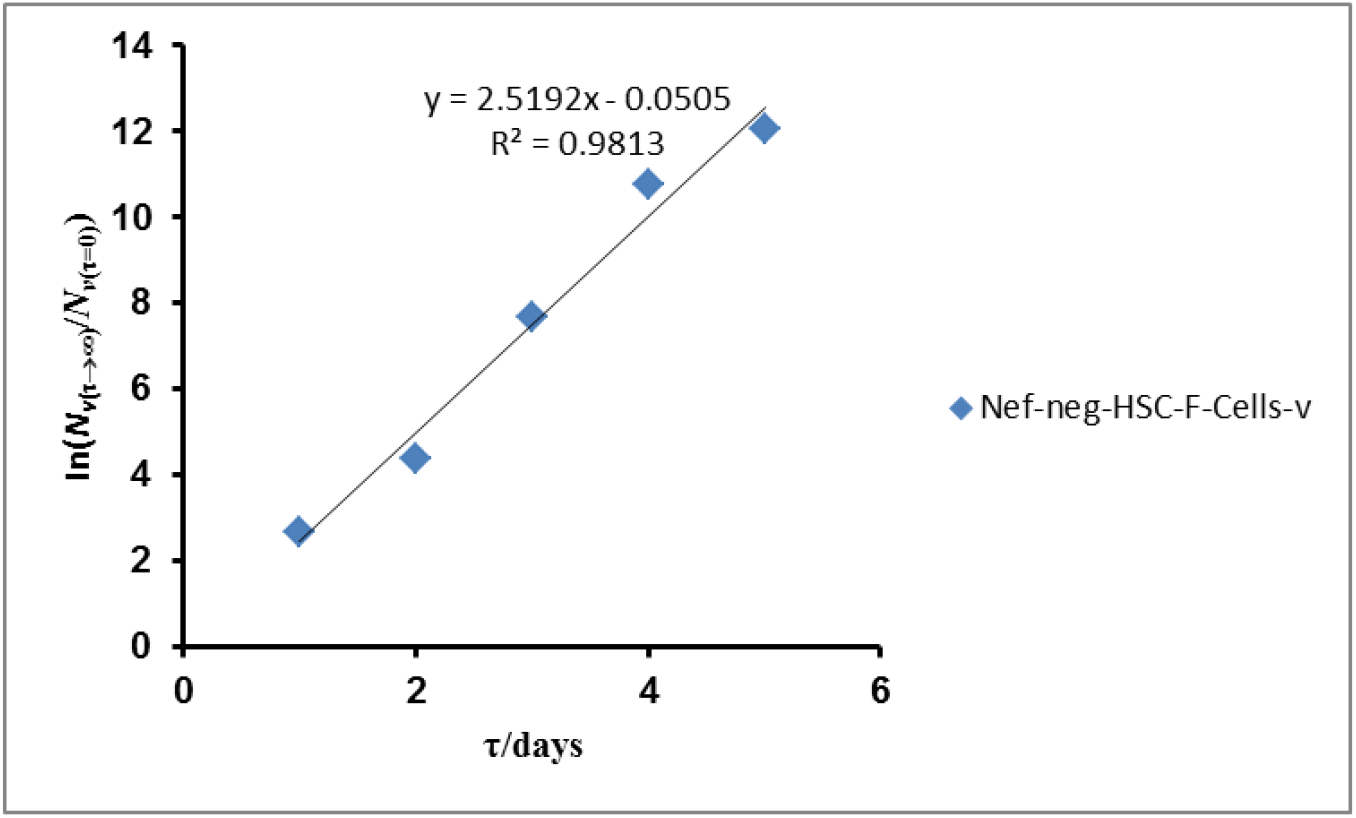
Rate constant for the replication of virus applied to Nef-NEGATIVE HSC-F CELLS (Nef-neg-HSC-F-Cells). The original data are as in previous reference.

**Figure 14:**
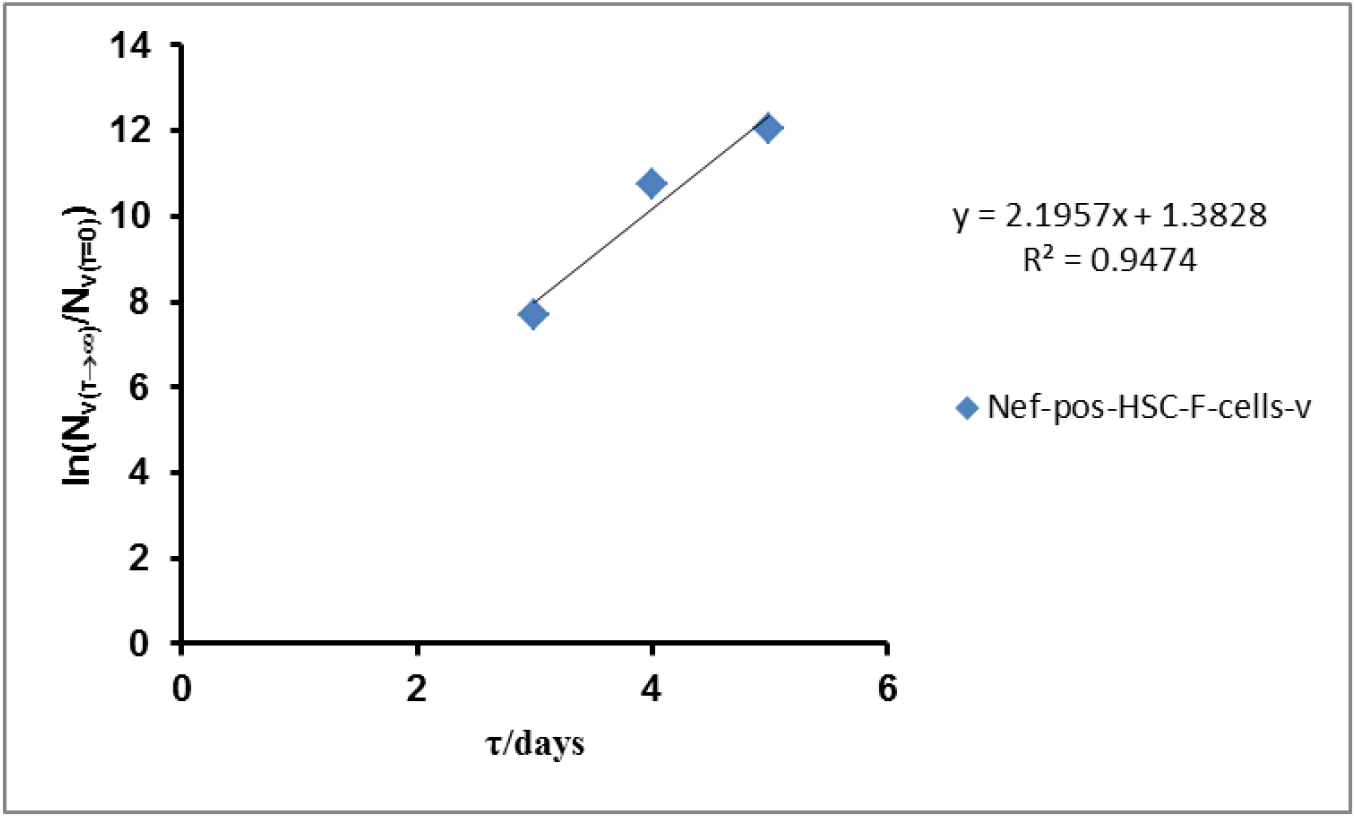
Rate constant for the replication of virus applied to Nef-POSITIVE HSC-F CELLS (Nef-pos-HSC-F-Cells). The original data are as in previous reference.

### 4.6. Time before symptom onset for a scenario where there are different viral populations with time beginning from the population in time = zero

As stated earlier, where there are different viral populations or loads on the increase as well as decreasing host cell population, a graphical approach is suitable for the earlier time lines before symptom onset. The data explored are available in the literature (Iwami *et al*., 2012). The graphical approach is accomplished based on Eq. (42). This gave Figures (15) and (16) for Nef-NEGATIVE HSC-F CELLS and Nef-POSITIVE HSC-F CELLS respectively. This is however, preceded by the determination of earlier viral load per unit time per cell based on the equation given as: *N*_*V*(τ=i)_/ (*τ=i*)*e*.(*k*_*v*_ (*τ=i*))= β*N*_*LR*(τ=i)_. Figures (17) and (18) for Nef-NEGATIVE HSC-F CELLS and Nef-Positive HSC-F CELLS respectively accomplished this purpose.

**Figure 15:**
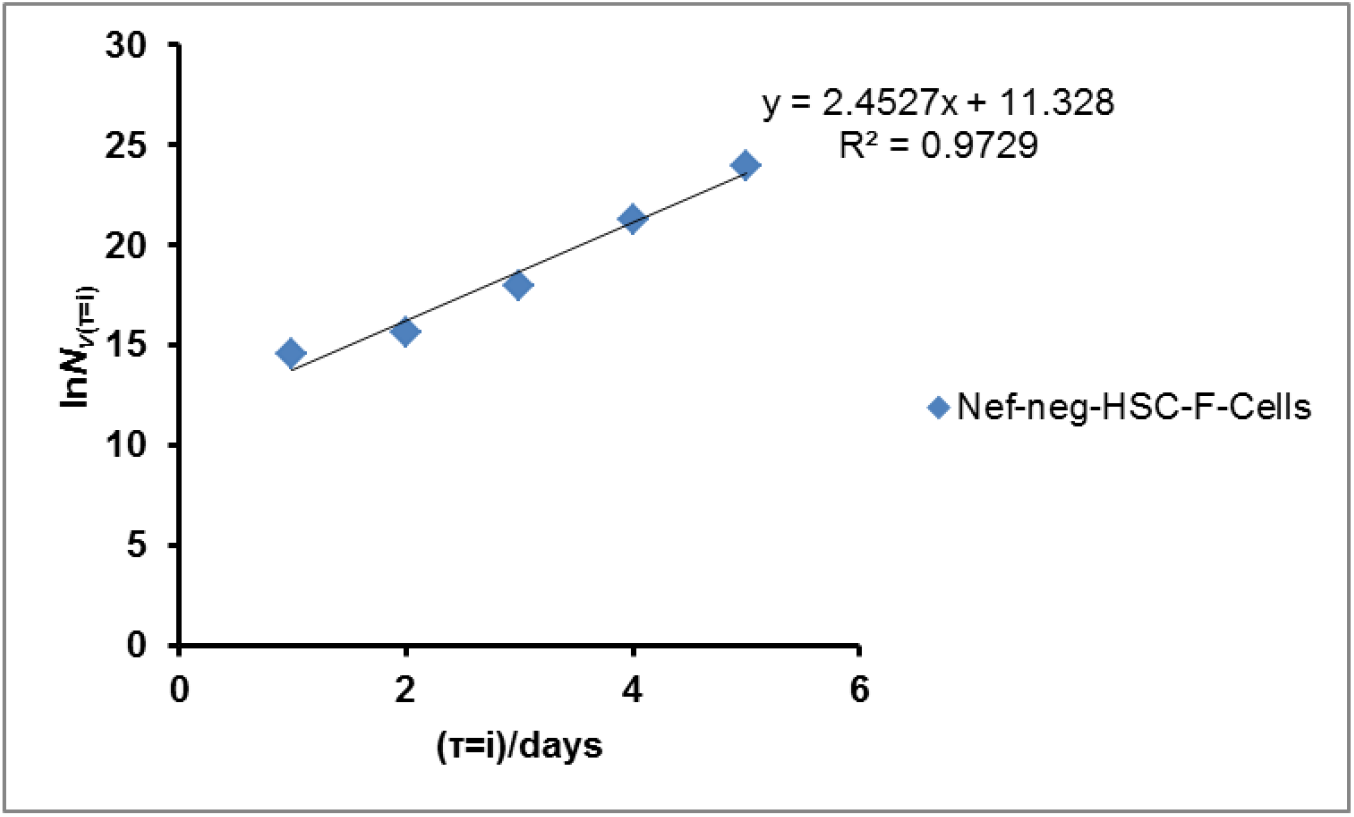
Determination of the earlier period of time before simptom onset with Nef-NEGATIVE HSC-F CELLS (Nef-neg-hs-f-cells). The original data are as in previous reference.

**Figure 16:**
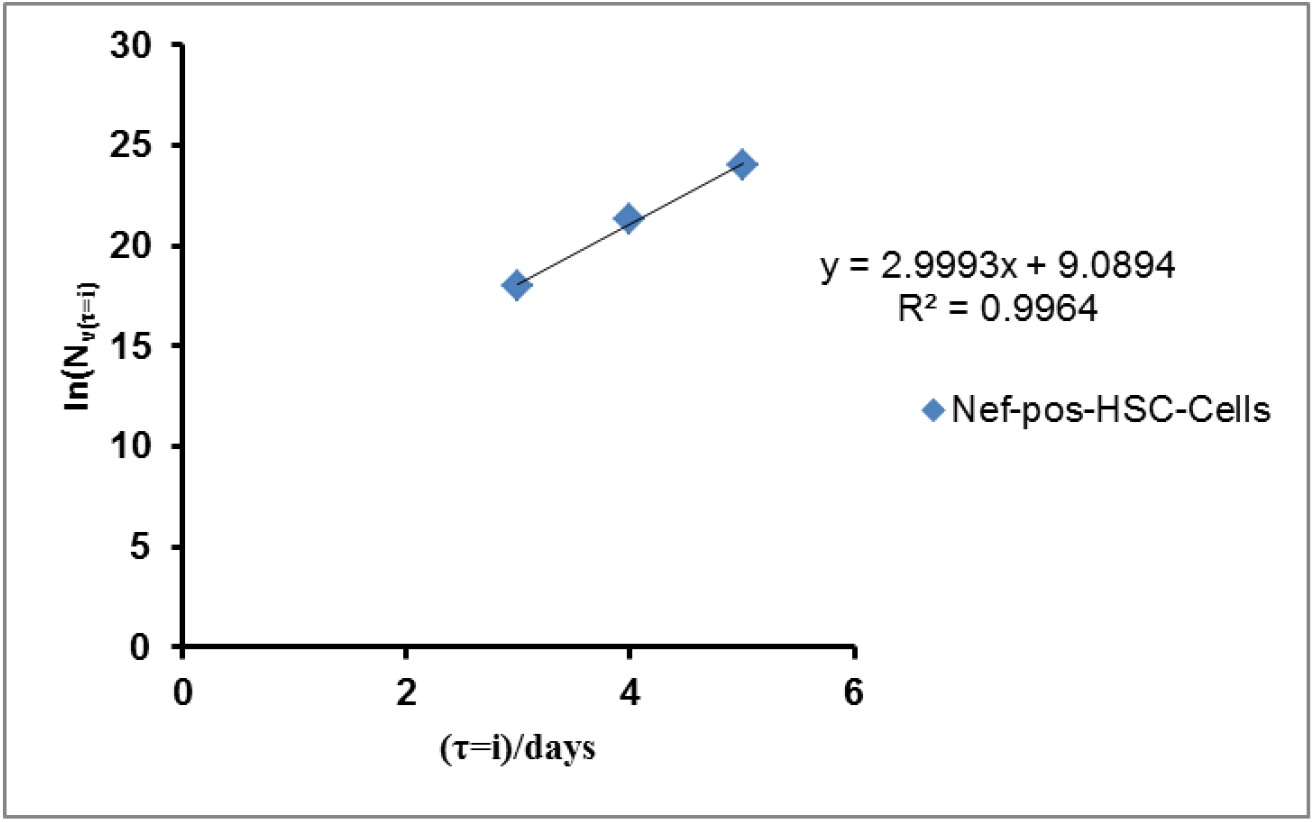
Determination of the earlier period of time before simptom onset with Nef-POSITIVE HSC-F CELLS (Nef-pos-hs-f-cells). The original data are as in previous reference.

**Figure 17:**
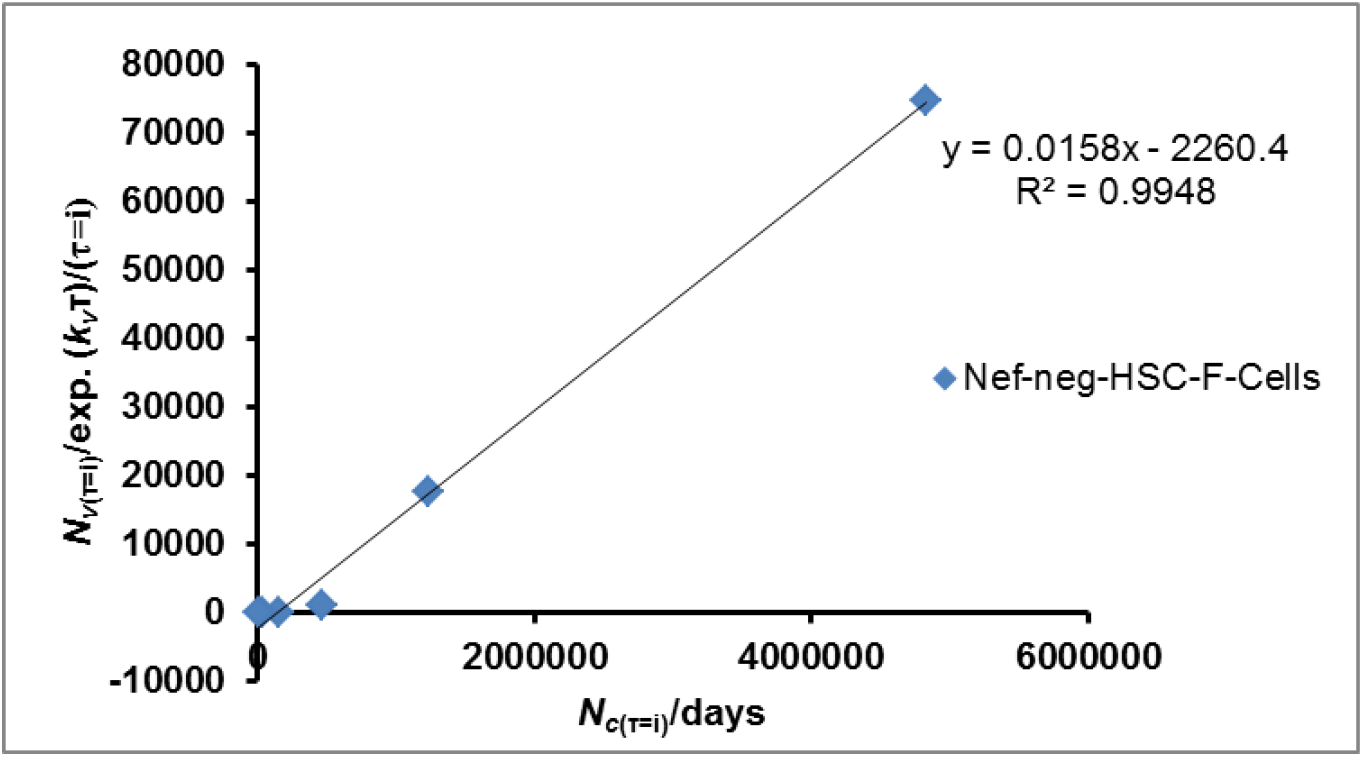
Determination of the number of virus per Nef-NEGATIVE HSC-F-CELLS per unit time (Nef-neg-hs-f-cells). The original data are as in previous reference. Period: 4-9 days.

**Figure 18:**
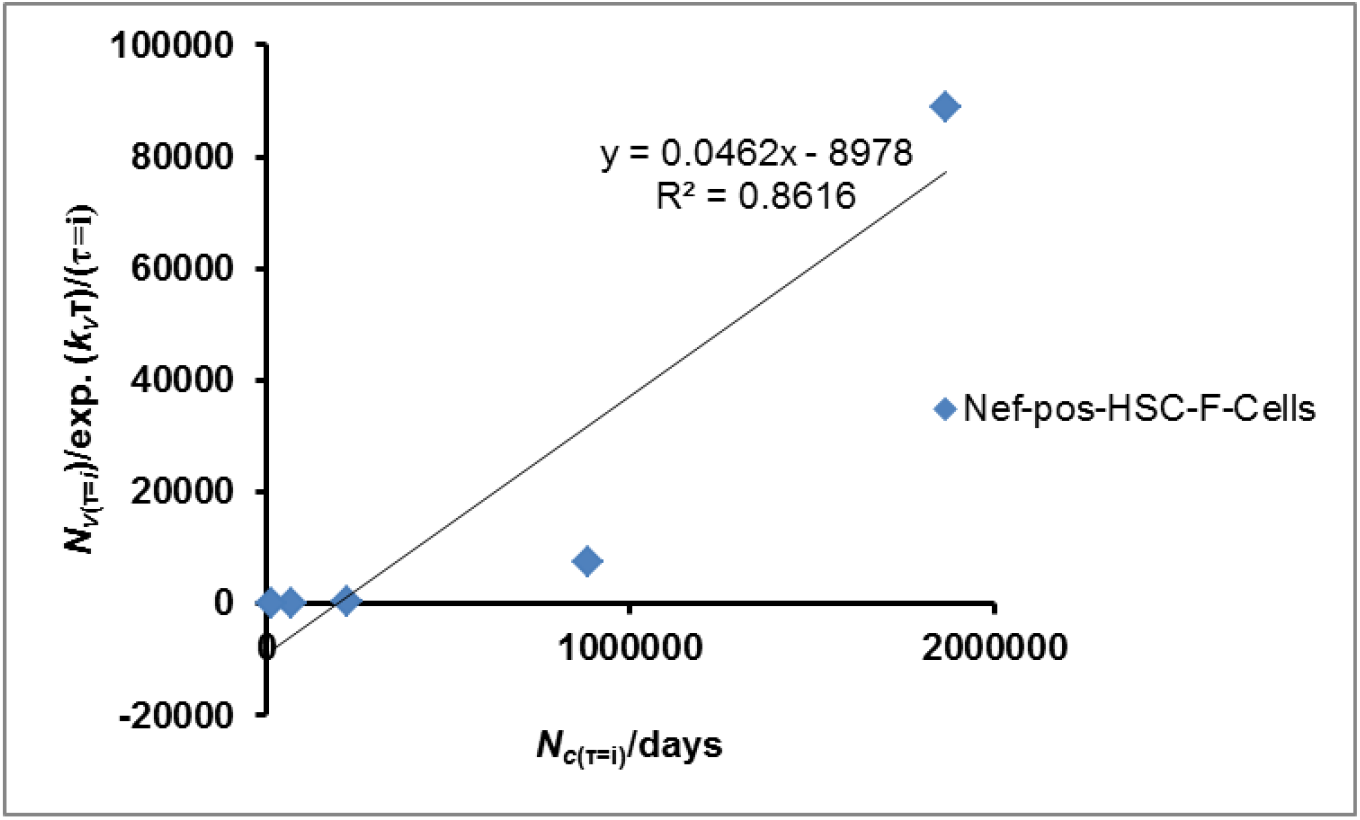
Determination of the number of virus per Nef-POSITIVE HSC-F CELLS per unit time (Nef-pos-hs-f-cells). The original data are as in previous reference. Period: 5-9 days.

### 4.7. Different times before symptom onset

There is duration (**τ**_***s***_) between the contacts made by the virus with the receptor of the target cell before symptom onset. The duration is determined by exploring the theoretically generated peak viral load (*N*_*v*(τ⟶¥)_) as time tends to infinity. This is based on the assumption that a virus with the highest rate constant of infection should have the highest viral peak load. Hence, the formula for the theoretical determination of peak viral load is given as: b_*x*_ *e*. (6)’ *e*. (5) / (β_1_ + β_2_ + β_3_), where *e*. (6) and *e*. (5) (Senders, *et al*., 2021) are, respectively, the number of infected cells and the number of viral particles (RNA copies to be specific) at any point in time; *b*_*x*_ is any of the rate constants of infection for different viruses namely, β_1_, β_2_, and β_3_. Although those values relate to the SARS-CoV-2 virus as indicated in the literature, they are generalized for the purpose of evaluating the equations.

The goal of exploring theoretically (Table 5a) and experimentally (Table 5c) generated viral load peaks at symptom onset is to reexamine the effectiveness of the methods and consistency in trend rather than in magnitude of the times computed. This concern is reflected in Tables 5b and 5d. In all cases, the relevant equations, Eqs (38), (39), (40), and (41) are suitable for single value of viral load theoretical and experimental at symptom onset and specific time at viral load peak, theoretical and experimental. This may not be the case if multiple viral peak values are given at different times. This aspect is addressed subsequently.

**Table 5a:** Theoretically determined viral load per ml expected from Eqs (38) and (39) using theoretically determined viral load peak.

| Virus | $N_{V(\tau \rightarrow \infty)}$ | Values based on Eq. (38) | Values based on Eq. (39) |
| --- | --- | --- | --- |
| **MERSCoV** | 2.66 e. (8) | 2.66 e. (-1) | 89.86 |
| **SARSCoV** | 9.31 e. (8) | 9.31 e. (-1) | 1.38 e. (-8) |
| **SARSCoV-2** | 9.88 e. (10) | 9.88 e. (1) | 1.11 e. (13) |

$N_{LR}(\tau=0) = e. (9)$ (Senders, *et al.*, 2021) and references therein (Iwami, *et al.*, 2012, Senders, *et al.*, 2021)

Sender *et al*. (2021) and references therein (Iwami, *et al*., 2012, Senders, *et al*., 2021) reports that the number of mucus cells in the nasal cavity is ∼ *e* (+9); this number is explored as the initial number of before infection.

**Table 5b:**
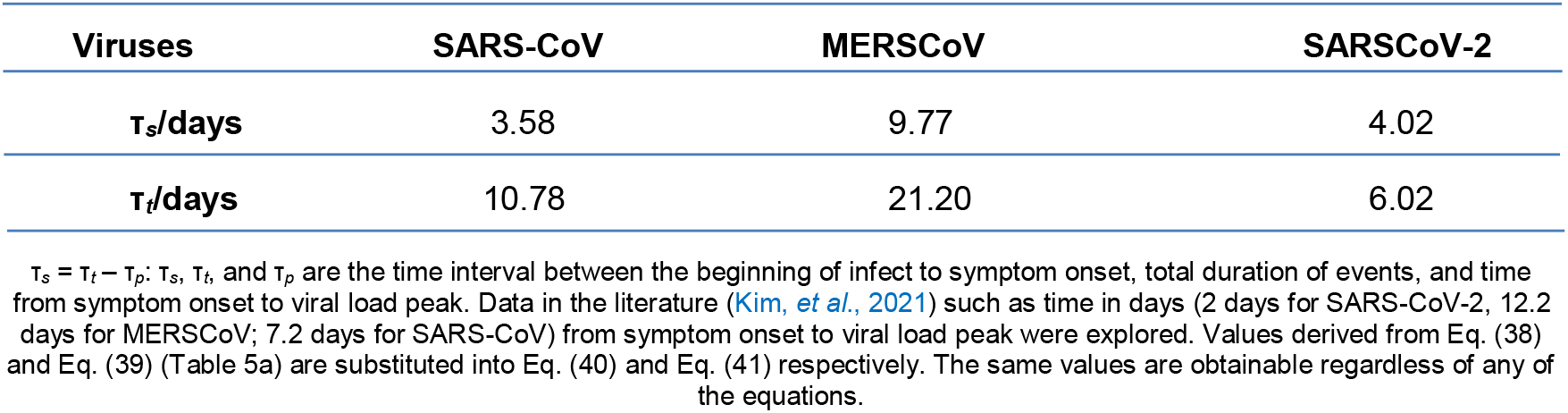
Time before symptom onset for events such as contact and binding to the cell membrane to the beginning of infections, symptom onset, and viral load peak.

**Table 5c:** Theoretically determined viral load per ml as expected from Eqs (38) and (39) at viral load peak determined based on experimental viral load at symptom onset.

| Virus | $N_{V(T \rightarrow \infty)}$ | Values based on Eq. (38) | Values based on Eq. (39) |
| --- | --- | --- | --- |
| **MERSCoV** | 3.95 e. (12) | 3.59 e. (3) | 1.21 e. (6) |
| **SARSCoV** | 2.71 e. (11) | 2.71 e. (2) | 4.02 e. (-6) |
| **SARSCoV-2** | 1.94 e. (7) | 1.94 e. (-2) | 2.18 e. (9) |

Experimental viral peak value of the virus is given as: $N_{V(T \rightarrow \infty)} = N_{V(T=0)} e. (k_v T)$ where, for the purpose of emphasis, $N_{V(T=0)}$ is the experimental viral load at symptom onset. Experimental viral load is sourced from Kim, *et al.* (2021)

Like the time lines (Table 5b) computed by exploring the theoretically generated viral load based on the theoretically computed viral load peak (Table 5a), the time lines (Table 5d) computed by exploring experimental viral load at symptom onset showed maximum value with MERS-CoV. Overall the values based on the experimental viral load at symptom onset are greater than those based on the theoretically determined viral load. Only the value for SARS-CoV-2 (Table 5d) based on experimental viral load at symptom onset is less than the value (Table 5b) computed based on theoretically determined viral load.

**Table 5d:**
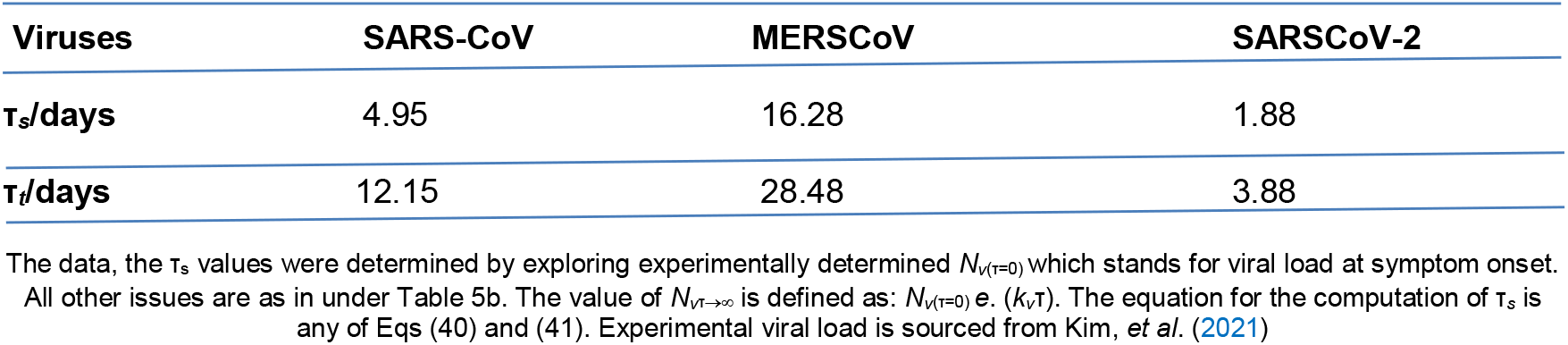
Determination of time before symptom onset using experimental *N*_*v*(τ=0)_ values for events such as contact and binding to the cell membrane to the beginning of infections, symptom onset, and viral load peak.

**Table 6:**
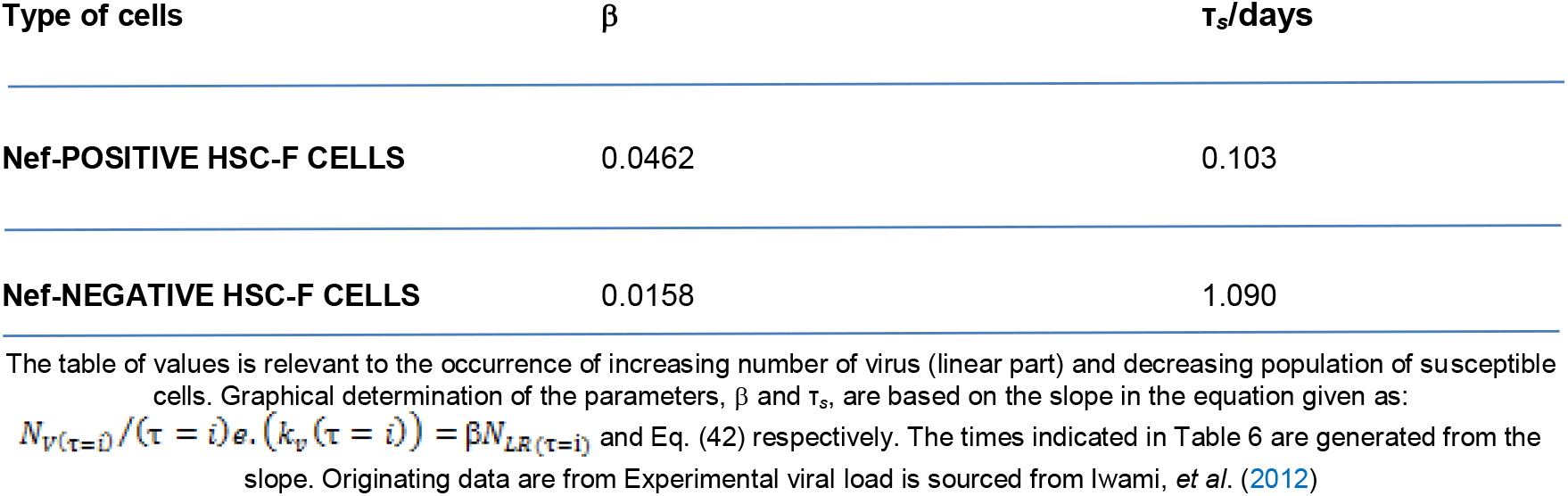
The viral load of shiv-ks661 (RNA copies/ml)/cell/unit time (β) and the time (τ_*s*_) interval between infection and earliest viral load in the presence of HSC-F CELLS.

| Type of cells | $\beta$ | $\tau_s/\text{days}$ |
| --- | --- | --- |
| Nef-POSITIVE HSC-F CELLS | 0.0462 | 0.103 |
| Nef-NEGATIVE HSC-F CELLS | 0.0158 | 1.090 |

The table of values is relevant to the occurrence of increasing number of virus (linear part) and decreasing population of susceptible cells. Graphical determination of the parameters, $\beta$ and $\tau_s$ , are based on the slope in the equation given as: $N_{V(\tau=i)}/(\tau=i) \text{ e. } (k_v(\tau=i)) = \beta N_{LR(\tau=i)}$ and Eq. (42) respectively. The times indicated in Table 6 are generated from the slope. Originating data are from Experimental viral load is sourced from Iwami, *et al.* (2012)

### 4.8. Variation of thermodynamic equilibrium constant for viral replication and cell death with time

This section which has been reported in a book submitted for assessment addresses the question of whether there is either upward or downward trajectory of equilibrium dissociation constant (*K*_*eq*(δ)_) with time. Fitting Eq. (21a) to variables that must be computed given first-order rate constants allows for this to be achieved. In the early stages of infection and other repercussions, the unitless *K*_*eq*(δ)_ is larger, as Figure 19 for Nef-Negative HSC-F Cells and Figure (20) for Nef-Positive HSC-F Cells illustrate. A power law governs the plot of *K*_*eq*(δ)_ against time (τ) in earlier days. Therefore, it is important that the medical team respond sooner rather than later in order to prevent the catastrophe that plagued the previous outbreak as a result of the authorities’ complacency. This is in line with empirically supported view that higher drug efficacy and earlier treatment initiation are associated with better outcomes: 74 % target cells remained uninfected after the course of infection – when treatment was initiated 1 day after symptom onset and antiviral effectiveness was 90 % (Kim, *et al*. 2021). The current results should dispel any doubt about the need to identify multiple earlier times starting from viral host cell contact leading to other detrimental events prior to cell death. This is especially relevant to those who are fortunate enough to have access to highly advanced, cutting-edge facilities. *In vitro* experimental results for the earlier times before symptom onset as outlined in Tables (5b), (5d), and (6) in different settings could serve as guide in developing and applying appropriate antidotes against the virus.

**Figure 19:**
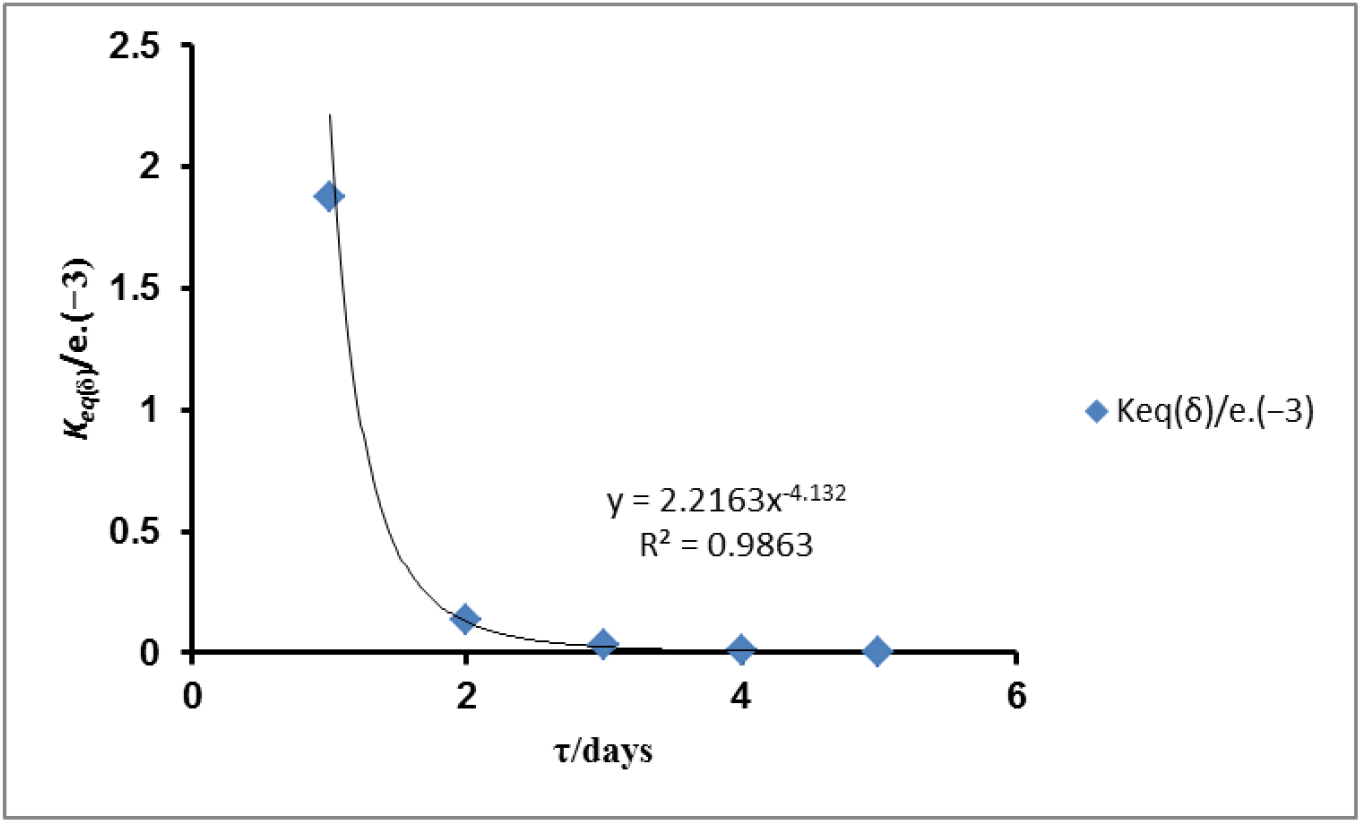
A plot of the dimensionless (unit less) equilibrium constant (*K*_*eq*(δ)_) for viral replication and cell death. The values of *k*_*LR*_ (rate constant for cell death) and *k*_*v*_ (rate constant for viral replication) are 1.1609 /day and 2.5192 /day respectively for Nef-Negative HSC-F Cells (Figures 11 and 12 respectively): Periods between 4 to 9 days were adopted. The day of detectable life Nef-Negative HSC-F Cells (a number = 6.4 *e*. (+6)) was on the zero day; the maximum number of cell was on the zero day while the initial viral load was 150096 at zero time; decreasing trend in the number of cells began from the 4^th^ to the 9^th^ day as reported by Iwami *et al*. (2012). The variation of *K*_*eq*(δ)_ with time (τ) in days obeys the power law.

**Figure 20:**
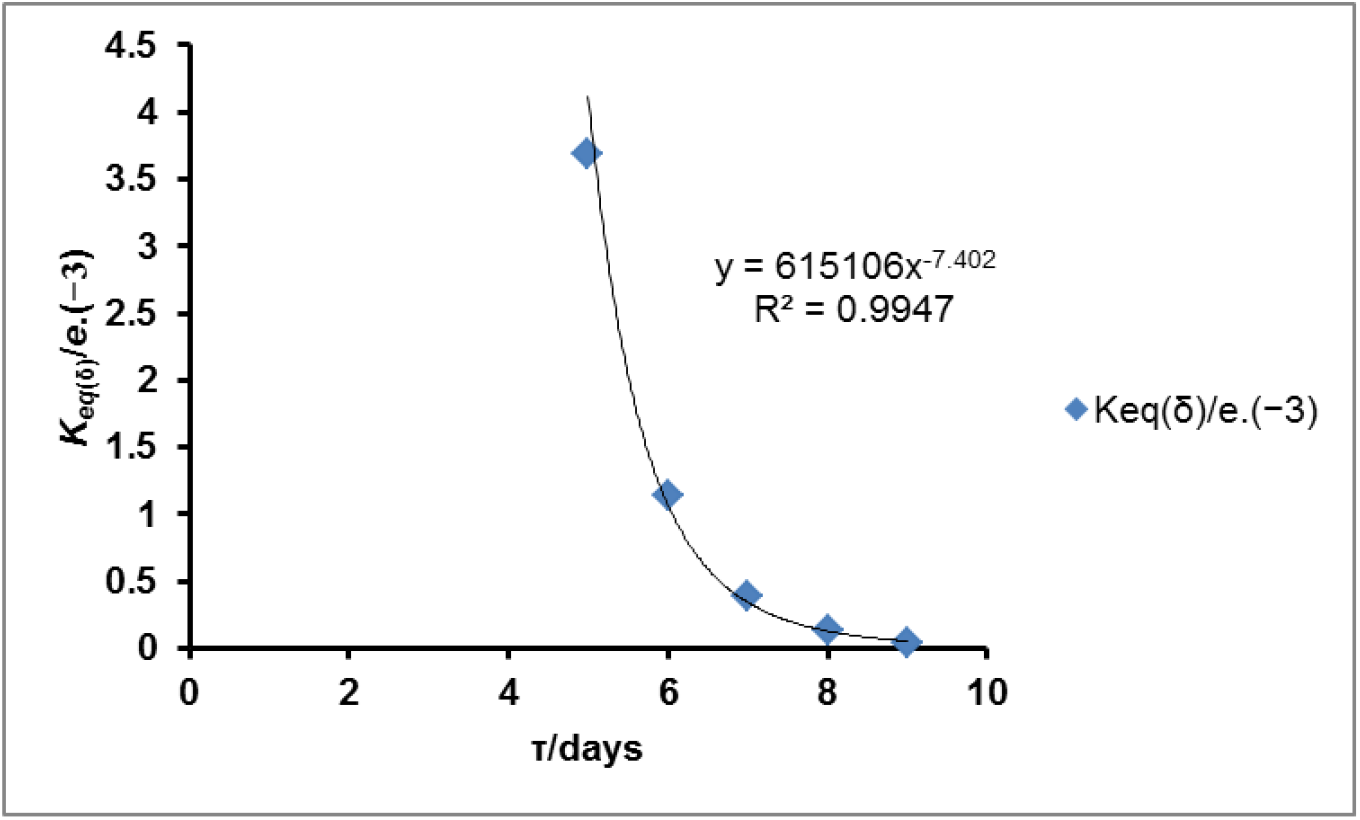
A plot of the dimensionless (unit less) equilibrium constant (*K*_*eq*(δ)_) for viral replication and cell death versus time. The values of *k*_*LR*_ (rate constant for cell death) and *k*_*v*_ (rate constant for viral replication) are 1.1241 /day and 2.1957 /day respectively for Nef-Positive HSC-F Cells (Figures 11 and 12 respectively): Periods between 5 to 9 days were adopted. The day of detectable life Nef-Positive HSC-F Cells (a number = 15392) was on the 3^rd^ day; the maximum number of cell was at the 5^th^ day while the initial viral load was 150096 at zero time; the decreasing trend in the number of cells began from the 5^th^ to the 8^th^ day as reported by Iwami *et al*. (2012). The variation of *K*_*eq*(δ)_ with time (τ) in days obeys the power law.

A virus of almost any kind is an obligate parasite since it needs a host for its multiplication; the host provides a suitable environment in terms of genetic, biochemical, and biophysical needs; the host becomes deprived of those suitable environments. Thus, this hypothesis considers, as a first step, the biophysical perspectives (made possible by the composition of the microbiome) on the implications of the complex environment of the cell that either enhance or inhibit the viral infection that seemed to have been taken for granted in the early days of the advent of COVID-19. The composition consists of proteins, lipoproteins, supra-biomolecules, lipids, including, most importantly, cholesterol, and a variety of biomolecules (some may be xenobiotics). These substances can all affect the microviscosity of the microbiome; this situation offers a favorable platform for the coronavirus to anchor itself, but it can also impede the spread of infectivity, suggesting that viscosity and cholesterol are two-edged swords in a pathophysiologic state orchestrated by SARS-CoV-2. This necessitates ideas or propositions (“hypotheses”) that can prevent viral infection and the related pathophysiology that leads to a high death rate.

### 4.9. Further biophysical consideration

The foci of this section are the impact of viscosity on translational motion of biomolecules that specifically target specific destination where they are needed. Entropic effects due to poly-dispersity also play an important role, altering the crowder mobility via two different mechanisms: 1), a more efficient crowder packing; and 2), an attractive contribution (known as depletion force (Crapo, *et al*., 1982) to the total force experienced by big molecules. All these increase the viscosity of the medium. “For diffusion to be observed in crowded environments without the formation of extensive aggregates, the intermolecular interactions must decrease compared to those derived in isolation” (Trovato and Tozzini, 2014). This has the potential to decrease the viscosity of the medium. The fact that the viscosity of a medium affects the rate of catalysis can be qualitatively and computationally evaluated given appropriate equations in this research. But first, the misinterpretation of such an equation is not unlikely, and so the equations are first analyzed in a manner that reflects the impact of viscosity on translational velocity that delivers biomolecules to the point of need.

In the equation such as :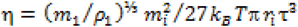, the product η*r*_*i*_τ^3^ is directly proportional to 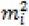, whereas ητ^3^ is inversely proportional to the Kelvin temperature. This seems to validate the claim that large molar mass solutes, either as substrates or viscogenes, have a greater impact on viscosity. One cannot ignore the fact that even a very high homogenous concentration of gelatinized starch, for instance, retards, if not halts, the mobility of the molecules because of hyperviscosity. In Eq. (5), the time (τ) taken to attain uniform concentration or terminal velocity is directly proportional to the square root of the product of RMSD and the mass of the solute and inversely proportional to the square root of Kelvin temperature. These cannot be unreasonable.

Again, high cholesterol concentrations—more especially, high concentrations of protein and saturated lipids—have a significant impact on the development of high viscosity. The viscosity state can have an upward or downward effect on the translational and rotational kinetic energy (*E*_*k*_). However, the primary focus is on translational velocity and *E*_*k*_. The translational *E*_*k*_ and velocity affect the targeted delivery of biomolecules, xenobiotic-like drugs, *etc*., to the site of need. Equation (2) clarifies these arguments for the impact of viscosity. In it, *E*_*k*_ is jointly and directly proportional to the cube root of the squared product of Kelvin temperature and translational diffusion coefficient. Consequently, *E*_*k*_ should also be inversely proportional to the cube root of the squared viscosity coefficient. Also, the translational velocity (*u*_*trans*_) is directly proportional to the cube root of the translational diffusion coefficient; it is also inversely proportional to the cube root of the viscosity coefficient. All these clearly explain why the rate of biological activities can be retarded if the viscosity of the medium is high.

Next, there is a need to give quantitative evidence on the issues raised. First the equation of the instantaneous translational velocity (the velocity before attaining the lower terminal velocity) is given as :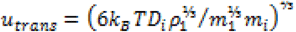 ; literature values of apparent diffusion coefficient of glucose in the cytosol and water at 310.15 K are 3.1 *e*. (−11) and 8.7 *e*. (−10) m^2^/s respectively (Kreft, *et al*., 2013). Fitting the equation above to each of the datum and molar mass of glucose (180 g/mol.) gives instantaneous velocity before terminal velocity as follows: ∼ 0.046674 m/s (cytosol); 0.141837 m/s (water).

The equation of RMSD is given as 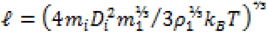. Fitting the equation to each of the datum gives: ∼ 1.328374 nm (cytosol); 12.267518 nm. The equation for thermochemical potential field force (TCPFF) is given as : 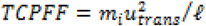. When the equation is fitted to the previously determined values of the parameters, the result is approximately 4.901357 e. (−19) N for water and cytosol. It is approximately equal to 3.0593 eV /m. The terminal velocity equation is as follows : 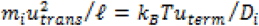. After solving for the terminal velocity (*u*_*term*_) and fitting the equation to the computed parameters and the molar mass of glucose, the results are approximately equal to 3.548614 nm /s for the cytosol and 99.590626 nm /s for the water. The magnitude of translational velocity and, ultimately, the pace of biogenesis and viral replication are determined by viscosity and its causal components, as is evident from the data thus far.

### 4.10. Hypothesis

Consequently, the following hypotheses may help halt the increase in viral load: 1) Increasing the mobility of PM’s microdomains, or lipid rafts, by reducing their cholesterol concentration; 2) adding dehydrating agents that are nano-encapsulated to the cytosol, where they become extremely hydrated at the expense of the solutes in order to increase the viscosity; 3) using nanoparticles or any other method that hasn’t been found yet to add cholesterol and neutral crowders, small and somewhat large, to the cytoplasm to increase the concentration of solutes of various sizes; and 4) looking into ways to make the cytoplasm and endoplasmic reticulum produce less cholesterol by raising the *K*_*M*_ and/or *K*_*d*_.

The following studies from the literature support the concepts or suggestions mentioned above: Although it is known that Cholesterol on cell membranes promotes viral entry, the significance of intracellular Cholesterol in SARS-CoV-2 infection cannot be overstated: The genes (from CRISPR libraries) necessary for cholesterol metabolism in SARS-CoV-2 infection include membrane-bound transcription factor peptidases, sites 1 and 2 (MBTPS 1 and MBTPS 2), low-density lipoprotein receptor (LDLR), sterol-regulatory element-binding protein (SREBP-2), and SREBP cleavage-activating protein (SCAP), as appropriate at the membrane level. As a result, lowering the amount of cholesterol in the membrane may require the use of peptidase inhibitors (MBTPS 1 and MBTPS 2). Additionally, amlodipine, a calcium ion channel antagonist, raises intracellular cholesterol levels, which may increase cytoplasmic viscosity and thus significantly reduce SARS-CoV-2 infection (Zhang, *et al*., 2020).

According to research, enhancing HDL’s cholesterol efflux capacity may lessen the severity of COVID-19 by influencing the ACE2 receptor’s translocation to lipid rafts, which includes cholesterol depletion of lipid rafts, and thereby the virus’s ability to infiltrate cells (Stadler, *et al*., 2022). Moreover, after methyl-β-cyclodextrin treatment, the level of ACE2 and furin protease in lipid rafts decreased due to the observed removal of plasma membrane cholesterol. However, when cells were loaded with cholesterol through treatment with apolipoprotein (Apo) E and serum, ACE2 and furin protease were trafficked to the lipid rafts, leading to an increase in SARS-CoV-2 infection (Wang, *et al*., 2021). Scavenger receptor B type I (SR-BI), commonly known as the HDL receptor, aids in the maintenance of cellular cholesterol homeostasis by promoting the selective uptake of cholesterol esters from HDL particles. HDL may drain cholesterol from the cell membrane, but it may also deposit cholesterol in the cytoplasm as part of its export function (Saddar *et al*., 2013). The SARS-CoV-2 S1 subunit attaches itself to apolipoprotein D and cholesterol as well as other HDL components before fusing. Consequently, reducing HDL particle concentrations may be a helpful therapeutic strategy to lessen SARS-CoV-2’s contagiousness.

### 4.11. SUMMARY

The cytoplasm and the membrane are both multicomponent media, but the membrane is more viscous, denser, and more compact than the cytoplasm. The biophysical perspective on the leveraging of the coronavirus’s anchoring and final infectivity that starts at the membrane level is based on these enabling physicochemical properties. When the organelles, especially the mitochondria, malfunction, tissue damage occurs first, followed by organ damage, system dysfunction, and premature death. The stability of the virus after initial contact and binding is improved by high viscosity in local microdomains called lipid rafts, which are enriched by high concentrations of cholesterol, phospholipids, sphingolipids, and various proteins like ACE2. This is done against the backdrop of a supportive asymmetric distribution of lipids in the two halves of the bilayer. These occurrences are always accompanied by activation energy and equilibrium thermodynamics. A suitable thermodynamic equilibrium constant that is dimensionless is required by the equilibrium thermodynamics component.

This study has been based on the dimensionless (unitless) equilibrium constant. Higher concentrations and viscosities of the causative chemicals, especially cholesterol, enhance infectivity at the membrane level even if they inhibit viral development and intercellular infection spread. Numerous medications can prevent the virus from sticking to the membrane, inhibit viral RNA replication, and increase the virus’s contagiousness. Unexpectedly high-pressure-control medications are among them. As evidenced by the decreasing trend in the unitless equilibrium constant with time, infectivity is most thermodynamically feasible at lower temperatures with lower activation energy; further development of infectivity is more possible during the early phase of infection (or prior to the onset of infection). The decreasing trend of the unitless equilibrium constant over time, which fits a power law, provides evidence for this.

## 5. CONCLUSION

The micro-anatomical (cell death and replication of viral RNA or the viral particles themselves) and biomolecular metabolic levels, model equations were derived for the computation and graphical determination of dimensionless equilibrium constants. The impact of the multi-complex components of the cytoplasm and cell membrane was described by the model that was developed and used. Based on the developed equations, the earlier times prior to the commencement of infection, peak viral load, and fall of susceptible cell population were ascertained graphically and computationally, where applicable. The results of the analysis of viral binding thermodynamics and activation energy revealed that the latter was more feasible at lower temperatures than at higher ones, while the former was lower. It was implied that the dimensionless constant values were higher at the earlier time of the infection and decreased with time, exhibiting a power law relationship. According to research findings and the calculated instantaneous and terminal velocities, viscosity and its causative compositional factors—cholesterol in particular—were found to have a dual-edged effect on the pathophysiologic state orchestrated by SARS-CoV-2. The following antidotes were suggested in light of these observations. It may also be necessary to reduce dietary cholesterol sources; biologically neutral viscogenes are added to the cytoplasm to increase viscosity, and medications that decrease cholesterol levels in the lipid raft and raise them in the cytoplasm are used to decrease and increase viscosity, respectively. It is advised that pharmaceuticals (including airborne surfactants) and drugs in solution be given at temperatures above body temperature. The efflux function of HDL in regulating cholesterol levels should be reinforced. Swab testing should be performed on a regular basis to detect significant infections early. Future *in vitro* and *in vivo* studies on viral infection might focus on various time periods at various temperatures, both above and below body temperature.

## AUTHOR CONTRIBUTIONS

The sole author designed, analyzed, interpreted and prepared the manuscript.

## COMPETING INTERESTS DISCLAIMER

The sole author has declared that he has no known competing financial interests OR non-financial interests OR personal relationships that could have appeared to influence the work reported in this paper.

## DISCLAIMER (ARTIFICIAL INTELLIGENCE)

The sole author hereby declare that NO generative AI technologies such as Large Language Models (ChatGPT, COPILOT, etc.) and text-to-image generators have been used during the writing or editing of this manuscript.

## ACKNOWLEDGMENT

I am very grateful to my siblings for their financial and in-kind support.

## INFORMED CONSENT STATEMENT

Not applicable

## HUMAN AND ANIMAL RIGHTS STATEMENT

Not applicable

## DEDICATION

For their servant-based leadership without dividing boundaries, this study is dedicated to Professor Ambrose Folorunsho Alli (late professor of medicine), the former governor of the now-defunct Bendel State (which later became Edo and Delta states), and his deputy, Chief Demas Onoliobakpovba Akpore (late teacher/principal, a man of hospitality and entertainer). Everyone was given bursaries without corruption, and everyone—students, academic and nonacademic candidates, and staff—was given the highest priority when it came to educational opportunities. Due consideration was given to health needs.

